# CiliaCarta: an integrated and validated compendium of ciliary genes

**DOI:** 10.1101/123455

**Authors:** Teunis J. P. van Dam, Julie Kennedy, Robin van der Lee, Erik de Vrieze, Kirsten A. Wunderlich, Suzanne Rix, Gerard W. Dougherty, Nils J. Lambacher, Chunmei Li, Victor L. Jensen, Michel R. Leroux, Rim Hjeij, Nicola Horn, Yves Texier, Yasmin Wissinger, Jeroen van Reeuwijk, Gabrielle Wheway, Barbara Knapp, Jan F. Scheel, Brunella Franco, Dorus A. Mans, Erwin van Wijk, François Képès, Gisela G. Slaats, Grischa Toedt, Hannie Kremer, Heymut Omran, Katarzyna Szymanska, Konstantinos Koutroumpas, Marius Ueffing, Thanh-Minh T. Nguyen, Stef J.F. Letteboer, Machteld M. Oud, Sylvia E. C. van Beersum, Miriam Schmidts, Philip L. Beales, Qianhao Lu, Rachel H. Giles, Radek Szklarczyk, Robert B. Russell, Toby J. Gibson, Colin A. Johnson, Oliver E. Blacque, Uwe Wolfrum, Karsten Boldt, Ronald Roepman, Victor Hernandez-Hernandez, Martijn A. Huynen

**Author notes:** Current address: Theoretical Biology and Bioinformatics, Science faculty, Utrecht University, Padualaan 8, 3584 CH Utrecht, the Netherlands. Current address: Agilent Technologies Sales & Services GmbH & Co. KG, Hewlett-Packard-Str. 8, 76337 Waldbronn. Current address: Centre for Research in Biosciences, University of the West of England, Coldharbour Lane, Bristol, BS16 1QY, UK. Current address: Max Planck Institute for Chemistry, Department of Multiphase Chemistry, Mainz, Germany (Jan F. Scheel). Current address: Department of Clinical Genetics, Unit Clinical Genomics, Maastricht University Medical Centre, P.O. Box 616, 6200 MD Maastricht, the Netherlands. Corresponding authors Teunis J. P. van Dam, PhD Prof. Martijn A. Huynen, PhD.

## Abstract

The cilium is an essential organelle at the surface of most mammalian cells whose dysfunction causes a wide range of genetic diseases collectively called ciliopathies. The current rate at which new ciliopathy genes are identified suggests that many ciliary components remain undiscovered. We generated and rigorously analyzed genomic, proteomic, transcriptomic and evolutionary data and systematically integrated these using Bayesian statistics into a predictive score for ciliary function. This resulted in 285 candidate ciliary genes. We found experimental evidence of ciliary associations for 24 out of 36 analyzed candidate proteins. In addition, we show that OSCP1, which has previously been implicated in two distinct non-ciliary functions, causes a cilium dysfunction phenotype when depleted in zebrafish. The candidate list forms the basis of CiliaCarta, a comprehensive ciliary compendium covering 836 genes. The resource can be used to objectively prioritize candidate genes in whole exome or genome sequencing of ciliopathy patients and can be accessed at http://bioinformatics.bio.uu.nl/john/syscilia/ciliacarta/.

---

Cilia are microtubule-based organelles extending from the surface of most eukaryotic cells, serving critical functions in cell and fluid motility, as well as the transduction of a plethora of sensory and biochemical signals associated with developmental processes^1^. Cilium disruption leads to a wide range of human disorders, known as ciliopathies, characterized by defects in many different tissues and organs leading to symptoms such as cystic kidneys, blindness, bone malformation, nervous system defects and obesity. The cilium is a complex and highly organized structure, typically comprised of a ring of nine microtubule doublets extending from a centriole-derived basal body, and enveloped by an extension of the plasma membrane. Importantly, cilia are compartmentalized structures, with a membrane and internal composition that differs significantly from that of the plasma membrane and cytoplasm^2,3^. Several hundred proteins are thought to be involved in the formation and function of ciliary structures and associated signaling and transport pathways. Between Gene Ontology (GO)^4^ and the SYSCILIA Gold Standard (SCGS)^5^ about 600 genes have already been associated with ciliary function. However, it is likely that many more ciliary proteins remain to be functionally characterized, as new ciliary genes are uncovered on a regular basis, often in relation to cilia-related genetic disorders.

Although omics data sets provide a rich source of information to determine the proteins that constitute an organelle, they are inherently imperfect. They tend to be biased towards proteins with specific properties, or lack coverage and sensitivity. It is well established that large scale approaches often miss key players, for example proteomics fares better at finding intracellular versus extracellular or membrane proteins^6^. Combining data sets into a single resource that exploits their complementary nature is a logical step. The power of such an approach has been demonstrated for various cellular systems such as small RNA pathways^7^, the innate antiviral response ^8^, and, most notably, by MitoCarta^9^. The latter is a compendium of mitochondrial proteins based on the integration of various types of genomics data that has been extensively used by the biomedical community^10^.

Like the mitochondrion, the cilium has been subject to approaches that exploit signals in genomics data to predict new ciliary genes, e.g. the specific occurrence of genes in species with a cilium^11,12^, the sharing of specific transcription factor binding sites like the X-box motif^13^ and the spatiotemporal co-expression of ciliary genes^14^. Furthermore, an existing database, CilDB, contains data from individual genomics, transcriptomics, and proteomics experiments related to the cilium^15^. Although such databases are invaluable to the researcher, it is not obvious how to handle the sometimes-conflicting information presented by independent data sources: i.e. how to weigh the data relative to each other. A powerful solution lies in statistical and probabilistic integration of data sets. Multiple methods exist to combine genomics data, ranging from simply taking the genes that are predicted by most data sources, to machine learning methods that take into account non-linear combinations of the data (reviewed in^16^). In this spectrum, naive Bayesian classifiers take a middle ground. They exploit the relative strengths of the various data sets while maintaining transparency of the integration. For each gene, the contribution of each data set to the prediction can be determined, and new, independent data sources can then simply be added to those already used to improve predictive value and coverage.

Here, we present CiliaCarta, an experimentally benchmarked compendium of ciliary genes based on literature, annotation, and genome-wide Bayesian integration of a wide range of experimental data, including a recent large-scale protein-protein interaction data set specifically focused on ciliary proteins^17^. We extend the currently known set of cilia-related genes with 228 putative cilia-related genes to a total of 836 genes. Based on the outcome of the probabilistic integration and the results from the experimental validations, we estimate the total size of the human ciliome to be approximately 1200 genes. Furthermore, we show that objective data integration using Bayesian probabilities is capable of overcoming biases based on other sources or literature. As an exemplar of our approach, we show that OSCP1, which several publications have described as a solute carrier or tumor suppressor, is also a ciliary component, shedding new light on previous observations.

## Results

### Data collection and curation

We collected and constructed a total of six new data sets from proteomics, genomics, expression and evolutionary data and complemented these with two public data sets (Table 1). The data sets include three new protein-protein interaction (PPI) data sets based on three methods: tandem-affinity purification coupled to mass spectrometry (TAP-MS)^17^, stable isotope labeling of amino acids in cell culture (SILAC), and yeast two-hybrid (Y2H) screens. The latter two data sets are published as part of this work and include 1666 proteins (1301 and 365, resp.) describing 4659 interactions (4160 and 499, resp.) and an estimated positive predictive value of X. Because the TAP-MS and SILAC data sets are based on similar methodology and have a large bait overlap (14 out of 16 SILAC baits were used in the TAP-MS data set), we merged them into a single data set (Mass-spec based PPI) to avoid bias (see methods). In addition, we created three bioinformatic data sets: (i)a data set describing the presence or absence of conserved cilia-specific RFX and FOXJ1 transcription factor binding sites (TFBS) in the promoter regions of human genes based on the 29 mammals project^18^, (ii) a gene expression screen for genes that co-express with a set of known ciliary genes, and (iii) a comprehensive co-evolution data set from the presence-absence correlation patterns of genes with the presence of the cilium over a representative data set of eukaryotic species. Supplementing these six data sets we included two published large-scale data sets, covering primary cilia and motile cilia: (i) the Liu *et al*. data set is a proteomics data set describing the protein content of sensory cilia derived from isolated murine photoreceptor cells^19^;the Ross *et al*. data set is a dynamic gene expression data set describing the up-regulation of genes during ciliogenesis in a time series after shearing off cilia in human lung epithelial cells^20^.

**Table 1:**
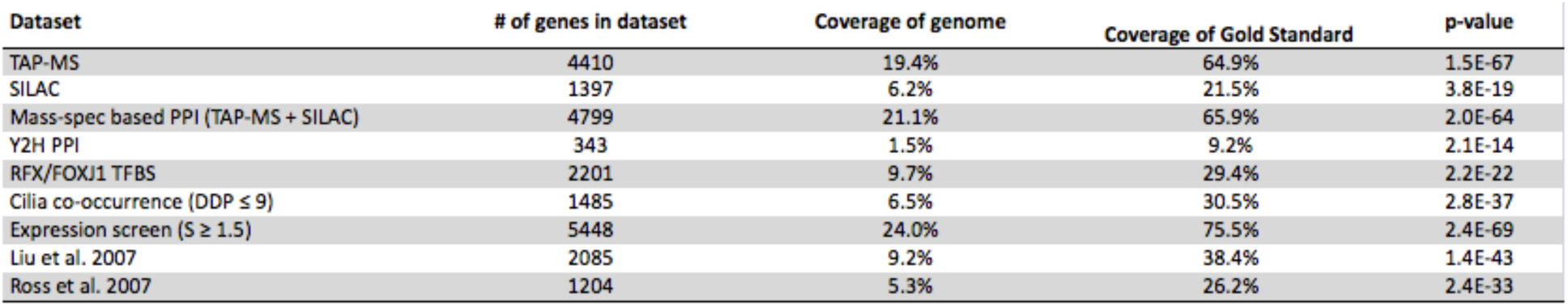
Coverage and predictive power of the cilium data sets. Coverage columns denote the fraction of the genome or SCGS that are actually identified by each approach. P-values indicate significant overrepresentation of the SCGS compared to random by Fisher’s exact test. Mass-spec based PPI represents the union of the TAP and SILAC data sets, resulting in the integration of five new and two published data sets.

| Dataset | # of genes in dataset | Coverage of genome | Coverage of Gold Standard | p-value |
| --- | --- | --- | --- | --- |
| TAP-MS | 4410 | 19.4% | 64.9% | 1.5E-67 |
| SILAC | 1397 | 6.2% | 21.5% | 3.8E-19 |
| Mass-spec based PPI (TAP-MS + SILAC) | 4799 | 21.1% | 65.9% | 2.0E-64 |
| Y2H PPI | 343 | 1.5% | 9.2% | 2.1E-14 |
| RFX/FOXJ1 TFBS | 2201 | 9.7% | 29.4% | 2.2E-22 |
| Cilia co-occurrence ($DDP \leq 9$) | 1485 | 6.5% | 30.5% | 2.8E-37 |
| Expression screen ($S \geq 1.5$) | 5448 | 24.0% | 75.5% | 2.4E-69 |
| Liu et al. 2007 | 2085 | 9.2% | 38.4% | 1.4E-43 |
| Ross et al. 2007 | 1204 | 5.3% | 26.2% | 2.4E-33 |

Although both data sets are not among the strongest predictors for ciliary function, they are of good quality and, importantly, have a much higher coverage of the human proteome than other published datasets (Supplementary fig. 1). In addition, the eight data sets were selected to ensure independence of the types of evidence and comprehensive coverage of molecular signatures for ciliary genes (for more details see methods). Each data set contains a highly significant signal for the discovery of ciliary genes (Table 1, p-values ranging from 2.1E-14 to 2.4E-69). A full description on each data set is given in the methods section.

### Bayesian integration of omics data sets provides gene-specific probabilities for cilia involvement

By combining the complementary sources of evidence for cilium function, we can in principle, obtain a data-driven, objective, high-confidence compendium of ciliary genes. To integrate the data sets in a probabilistic manner we assessed their efficacy at predicting ciliary genes using the SCGS, a manually curated set of known ciliary genes (the ‘positive’ set)^5^, and a set of genes whose proteins show non-ciliary subcellular localization and are most likely not involved in ciliary function (the ‘negative’ set, see methods). We divided each data set into appropriate sub-categories that reflect increasing propensities to report ciliary genes (Fig. 1a). Then, for each data category we calculated the likelihood ratios of predicting ciliary genes versus predicting non-ciliary genes using Bayes’ theorem. The final log-summed likelihood ratio, which includes the prior expected probability to observe a ciliary gene in the genome, we here call the CiliaCarta Score. This score therefore represents the likelihood for a gene to be ciliary, based on all the data sets considered (see methods; CiliaCarta and individual data set scores can be found in Supplementary table 1). The integrated score readily distinguishes between the positive and negative set (Fig. 1b, p-value: 2.8e-85 Mann-Whitney U test). A ten fold cross-validation demonstrates that the results are highly robust, with an area under the curve of 0.86 (Supplementary Fig. 2). The top ranking genes are highly enriched for known ciliary genes (Fig. 1c). The Bayesian classifier outperforms any individual data set, while achieving genome-wide coverage (Fig. 1d).

**Figure 1:**
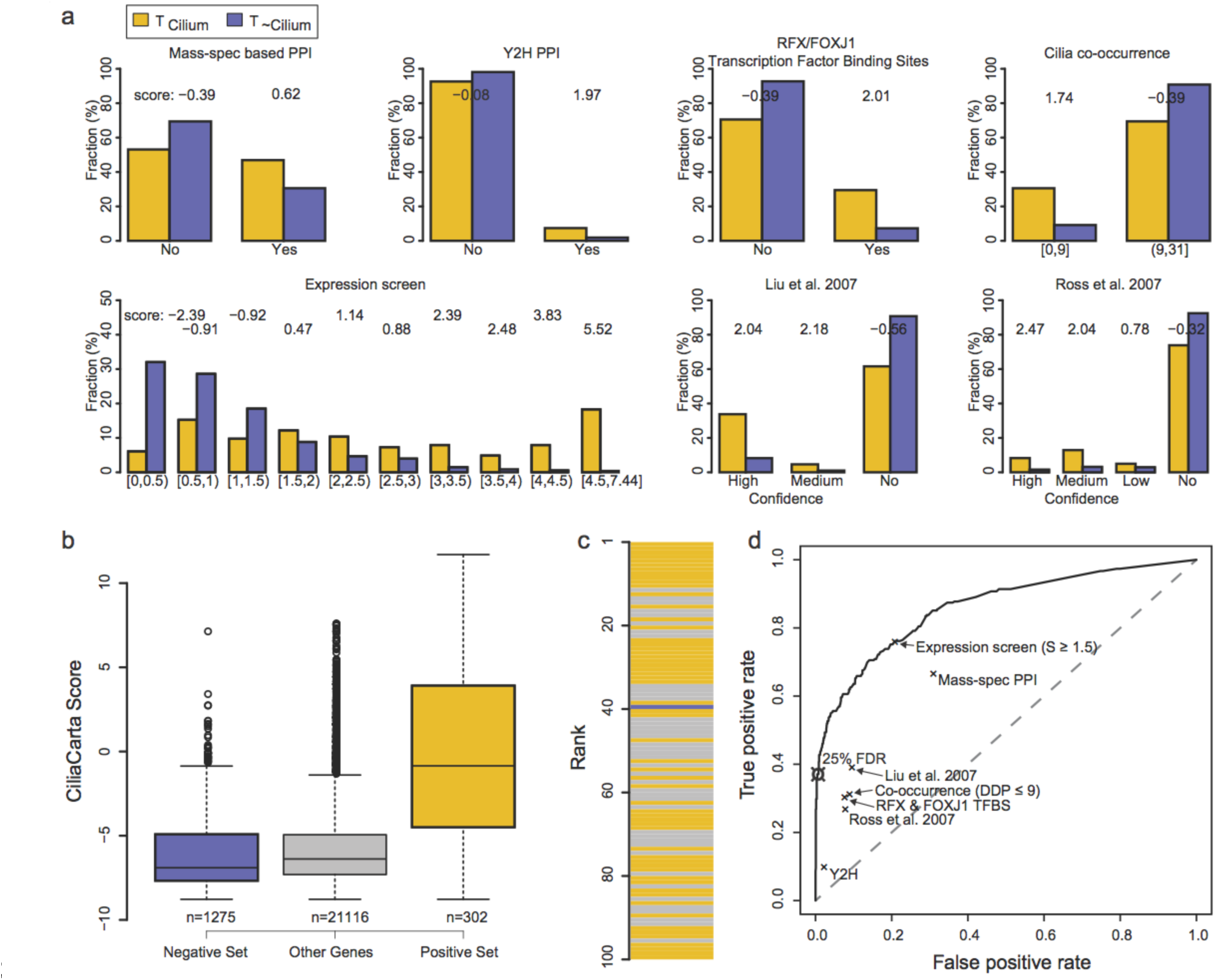
Data sets and performance of the Bayesian classifier for predicting ciliary genes. a) For each data set the fraction of positive (Tcilium) and negative gene sets (Tãcilium) and the log-likelihood scores are displayed per sub-category. b) Distributions of the integrated CiliaCarta scores for the negative set, the positive set, and the remaining unassigned genes. The positive set has significantly higher scores than the negative set (p-value: 2.8e-85 Mann-Whitney U test). c) Top 100 scoring genes. Known ciliary genes from the positive set are in yellow, genes from the negative set are in blue. High scoring genes in grey are prime candidate novel ciliary genes. d) Receiver-Operator Characteristics curve showing the performance of the Bayesian classifier as a function of the CiliaCarta Score and the performance of the individual data sets.

### Experiments validate ciliary function for 67% of selected candidate genes

In order to validate the quality of our approach we performed a series of experimental tests for ciliary function or localization of newly predicted candidate genes and their proteins. We set our inclusion threshold at a False Discovery Rate of 25% (cFDR, corrected for the prior expectation to observe a cilium gene, Methods). 404 genes fall within this threshold, 285 of which were not in the training sets and thus constitute novel candidate ciliary genes. Eight genes from the negative set (HSPH1, CALM3, COL21A1, CALM1, PTGES3, HSPD1, COL28A1, and PPOX) occur in the top 404 genes. Literature searches revealed no previous connection to the cilium. Some high scoring non-ciliary genes are expected due to stochasticity in the underlying experimental data, but it is possible that some of these genes will turn out to have a ciliary function in the future. The FDR for the 285 candidate genes is 33% when excluding genes from the training sets (Methods). From the candidates we selected a total of 36 genes, spread evenly across the top predictions, for experimental validation that were not known to be ciliary at the time (Table 2). The gene selection was performed as unbiased as possible and was restricted by orthology (zebrafish, *C. elegans*) and the resources available to the participating labs (see methods). We performed validations in human, mouse, zebrafish and *C. elegans* by applying six distinct approaches to determine ciliary localization and function.

**Table 2:**
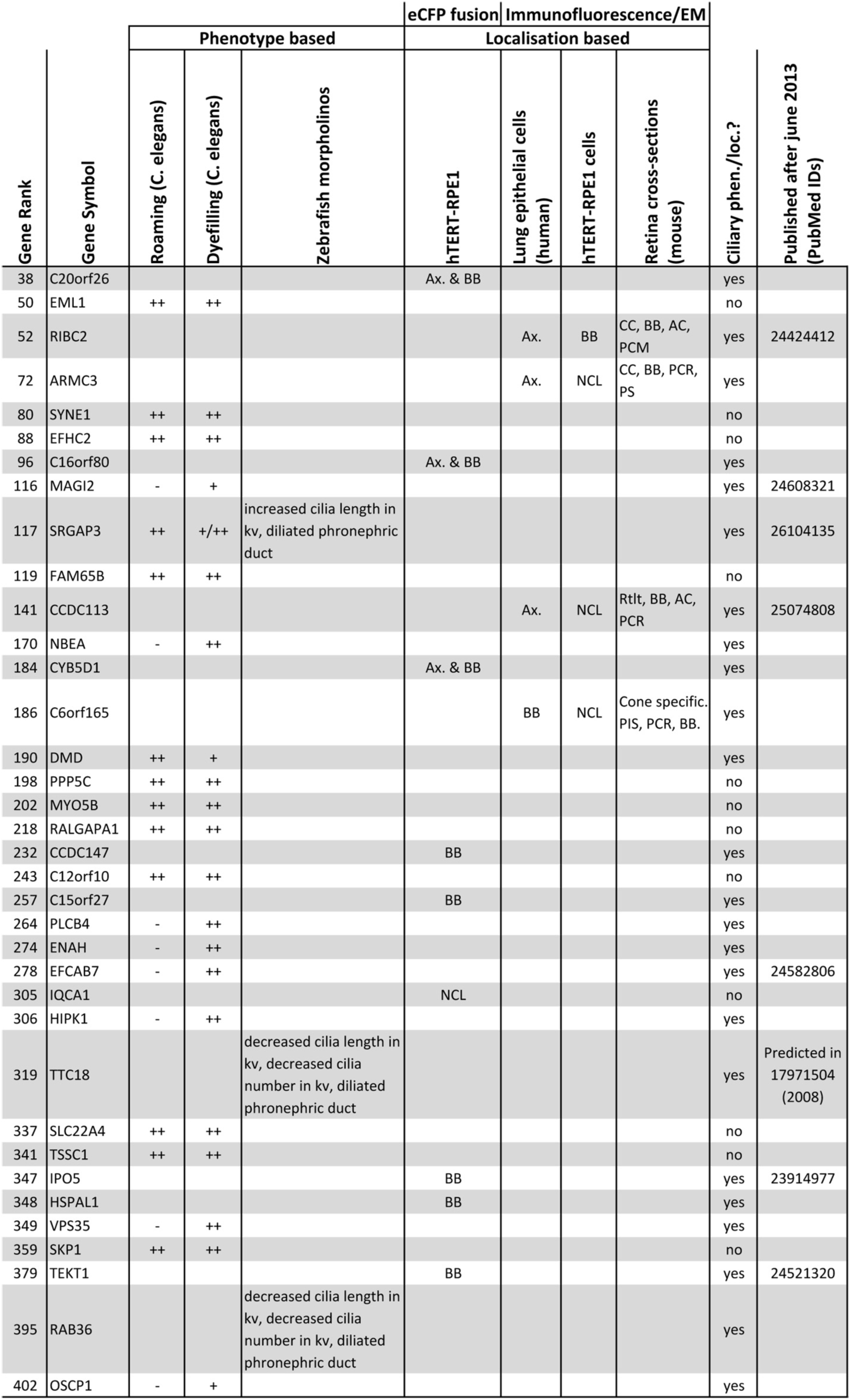
Genes selected for validation and the validation outcomes. Empty cells means not tested. Seven proteins (marked by *) have been published as ciliary proteins since the start of the validation experiments. Ax.: axonemal, BB: basal body, NCL: non-ciliary localization, CC: connecting cilium, AC: adjacent centriole, PCM: periciliary membrane, PCR: periciliary region, PS: photoreceptor synapse, Rtlt: Rootlet, PIS: photoreceptor inner segment. Roaming: “++” normal, defective (p<0.000l). Dye uptake: "++” normal, “*+*/*++*” slightly reduced uptake, “*+*” mild reduced dye uptake, “-” defective dye uptake.

| Gene Rank | Gene Symbol | Phenotype based |  |  | eCFP fusion | Immunofluorescence/EM |  |  | Ciliary phen./loc.? | Published after June 2013 (PubMed IDs) |
| --- | --- | --- | --- | --- | --- | --- | --- | --- | --- | --- |
|  |  | Roaming (C. elegans) | Dyefilling (C. elegans) | Zebrafish morpholinos | hTERT-RPE1 | Lung epithelial cells (human) | hTERT-RPE1 cells | Retina cross-sections (mouse) |  |  |
| 38 | C20orf26 |  |  |  | Ax. & BB |  |  |  | yes |  |
| 50 | EML1 | ++ | ++ |  |  |  |  |  | no |  |
| 52 | RIBC2 |  |  |  |  | Ax. | BB | CC, BB, AC, PCM | yes | 24424412 |
| 72 | ARMC3 |  |  |  |  | Ax. | NCL | CC, BB, PCR, PS | yes |  |
| 80 | SYNE1 | ++ | ++ |  |  |  |  |  | no |  |
| 88 | EFHC2 | ++ | ++ |  |  |  |  |  | no |  |
| 96 | C16orf80 |  |  |  | Ax. & BB |  |  |  | yes |  |
| 116 | MAGI2 | - | + |  |  |  |  |  | yes | 24608321 |
| 117 | SRGAP3 | ++ | + / ++ | increased cilia length in kv, dilated pronephric duct |  |  |  |  | yes | 26104135 |
| 119 | FAM65B | ++ | ++ |  |  |  |  |  | no |  |
| 141 | CCDC113 |  |  |  |  | Ax. | NCL | Rtl, BB, AC, PCR | yes | 25074808 |
| 170 | NBEA | - | ++ |  |  |  |  |  | yes |  |
| 184 | CYB5D1 |  |  |  | Ax. & BB |  |  |  | yes |  |
| 186 | C6orf165 |  |  |  |  | BB | NCL | Cone specific. PIS, PCR, BB. | yes |  |
| 190 | DMD | ++ | + |  |  |  |  |  | yes |  |
| 198 | PPP5C | ++ | ++ |  |  |  |  |  | no |  |
| 202 | MYO5B | ++ | ++ |  |  |  |  |  | no |  |
| 218 | RALGAPA1 | ++ | ++ |  |  |  |  |  | no |  |
| 232 | CCDC147 |  |  |  | BB |  |  |  | yes |  |
| 243 | C12orf10 | ++ | ++ |  |  |  |  |  | no |  |
| 257 | C15orf27 |  |  |  | BB |  |  |  | yes |  |
| 264 | PLCB4 | - | ++ |  |  |  |  |  | yes |  |
| 274 | ENAH | - | ++ |  |  |  |  |  | yes |  |
| 278 | EFCAB7 | - | ++ |  |  |  |  |  | yes | 24582806 |
| 305 | IQCA1 |  |  |  | NCL |  |  |  | no |  |
| 306 | HIPK1 | - | ++ |  |  |  |  |  | yes |  |
| 319 | TTC18 |  |  | decreased cilia length in kv, decreased cilia number in kv, dilated pronephric duct |  |  |  |  | yes | Predicted in 17971504 (2008) |
| 337 | SLC22A4 | ++ | ++ |  |  |  |  |  | no |  |
| 341 | TSSC1 | ++ | ++ |  |  |  |  |  | no |  |
| 347 | IPO5 |  |  |  | BB |  |  |  | yes | 23914977 |
| 348 | HSPAL1 |  |  |  | BB |  |  |  | yes |  |
| 349 | VPS35 | - | ++ |  |  |  |  |  | yes |  |
| 359 | SKP1 | ++ | ++ |  |  |  |  |  | no |  |
| 379 | TEKT1 |  |  |  | BB |  |  |  | yes | 24521320 |
| 395 | RAB36 |  |  | decreased cilia length in kv, decreased cilia number in kv, dilated pronephric duct |  |  |  |  | yes |  |
| 402 | OSCP1 | - | + |  |  |  |  |  | yes |  |

#### Validation by phenotype

In *C. elegans* we investigated candidate gene associations with cilium formation and function *in vivo*.In the *C. elegans* hermaphrodite (959 cells), cilia are found on 60 sensory neurons, most of which are located in the head. These cilia are immotile, eight of which extend from the dendritic endings, forming sensory pores in the nematode cuticle^21^. 68 of the 285 candidate genes have one-to-one orthologs in *C. elegans*. The apparent low number of orthologs in *C. elegans* could have a number of reasons, one of which is the absence of motile cilia, which resulted in the loss of ciliary genes related to motility. For 21 of these genes, viable alleles were available from the *Caenorhabditis* Genetics Center (University of Minnesota, USA) (Supplementary table 2). The available alleles were all nonsense or deletion mutations and predicted to severely disrupt gene function. For each available mutant, we employed dye-filling assays - to indirectly assess the integrity of a subset of cilia (six amphid pairs and one phasmid pair) - foraging (roaming) and osmotic avoidance assays to investigate sensory behaviors^22,23^. Notably, none of the mutants showed an abnormal osmotic avoidance response, or a highly penetrant dye uptake phenotype (Supplementary table 2), implying that the associated genes are not involved in global regulation of cilium formation or function. In addition all worm mutants had normal physical appearance. Roaming defects were observed for nine mutants, three of which also displayed a mild defect in dye uptake (Fig. 2a & b, Supplementary Fig. 3, and Supplementary table 2).

**Figure 2:**
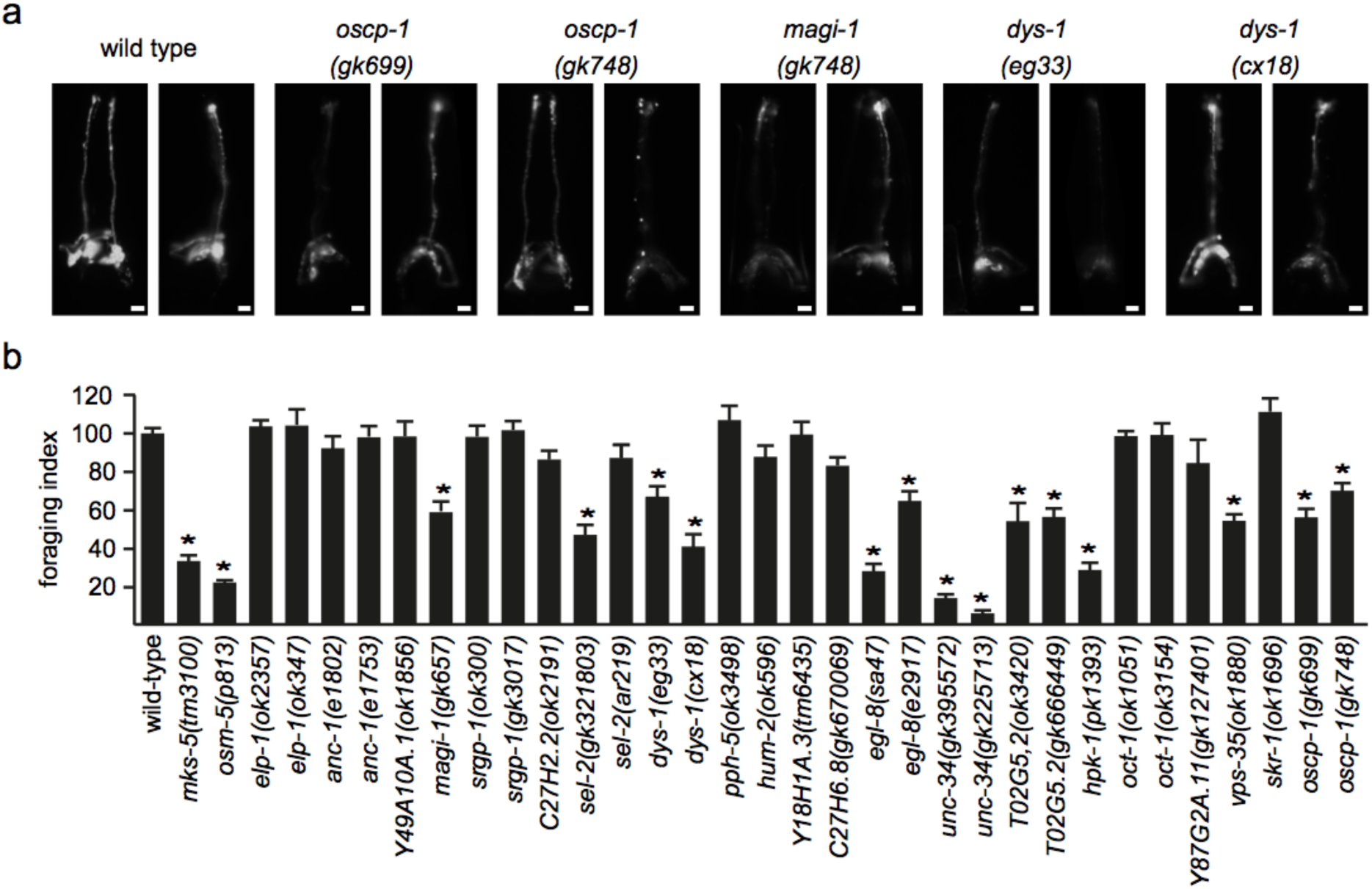
Validation by worm phenotype. a) *C. elegans* dye uptake assay. In wild type worms, DiI dye is taken up by 6 pairs of amphid (head) and one pair of phasmid (tail; not shown) neurons via their environmentally exposed sensory cilia. In the mutants shown, the amount of incorporated dye is modestly reduced, although most or all neurons still uptake the dye. Scale bars: 20 um. b) Single worm roaming assays. Bars represent mean ± S.E.M (n≥20) independent experiments), normalized to wild type control. * p<0.05 (unpaired t-test; vs. WT).

Three candidate genes with clear orthologs in zebrafish, *srgap3, ttc18* and *rab36* were investigated in zebrafish for cilia-related phenotypes after knockdown by morpholinos. The development of cilia in the Kupffer’s vesicle was examined (Fig. 3a). *rab36* and *ttc18* morphants show significantly reduced cilia length and number in the Kupffer’s vesicle (p<0.001 and p<0.05 resp.), while the *srgap3* morphant shows an increased cilia length (p<0.001), but no effect on cilia number. Pronephric ducts are several orders of magnitude larger in all three morphants during the pharyngula period (24 hpf) compared to wild type (Fig. 3b, p-value < 0.001). Furthermore, all three morphants exhibit body-axis defects associated with cilium dysfunction^24,25^ (Fig. 3c). Although, zebrafish morpholino phenotypes, and C. *elegans* phenotypes, do not provide definitive proof that these genes are all ciliary, they provide quantitative support for a substantial enrichment of ciliary genes in the CiliaCarta.

**Figure 3:**
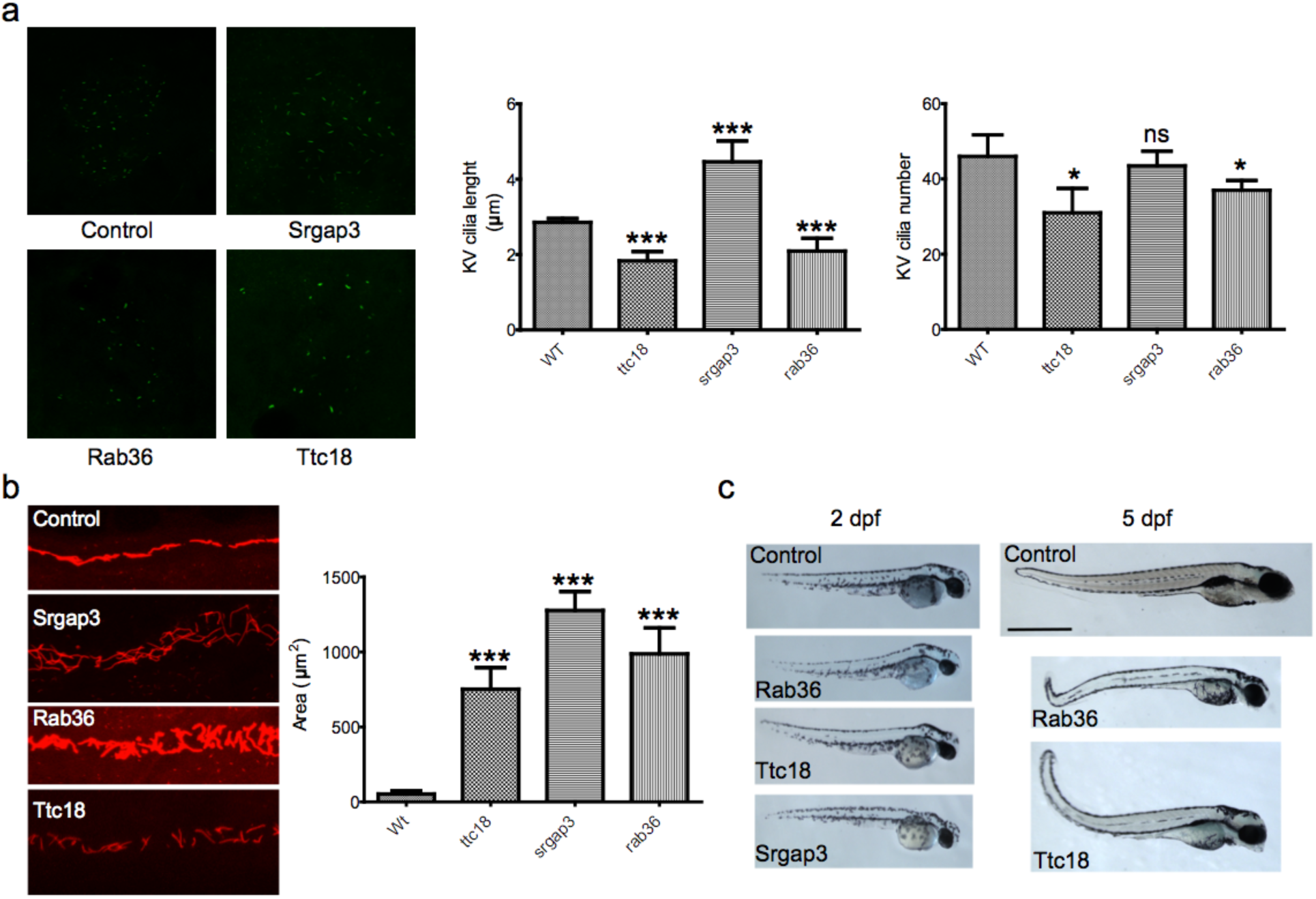
Validation by zebrafish phenotype. a) Cilia length and number in zebrafish Kupffer’s vesicles. The length and number of cilia in ttc18 and rab36 morphants are significantly reduced (p<0.001 and p<0.05 resp.). Cilia in srgap3 morphants are elongated (p<0.001) but the number of cilia is normal. Bars represent mean ± S.E.M. b) Pronephric ducts in 24 hpf morphants. Cilia are stained with antibodies against acetylated alpha tubulin. The pronephric ducts are significantly enlarged for all three morphants compared to wild type (p<0.001). Bars represent mean ± S.E.M. c) Whole embryo phenotype 2 days post fertilization (dpf) and 5 dpf zebrafish control and morphant embryos. All morphants exhibit the body curvature that is characteristic for cilia dysfunction. Note that in our screening we did not manage to obtain surviving srgap3 morphants past 3 dpf.

#### Validation by subcellular localization

Subcellular localization studies were performed for a total of nine proteins in ciliated hTERT-RPE1 cells using eCFP-tagged overexpression constructs (Table 2, Supplementary Fig.4). Eight proteins localized to the basal body and/or the axoneme. To account for possible localization artifacts, two representative photos are taken per eCFP fusion protein, containing at least one ciliated cell from the same slide. Cells transfected for c15orf22::eCFP and c16orf80 ciliated only when expression of the eCFP fusion protein was low. Fig. 4a shows representative examples, with C20orf26 and CCDC147 localizing to the basal body, whilst IQCA1 was not enriched at the cilium.

**Figure 4:**
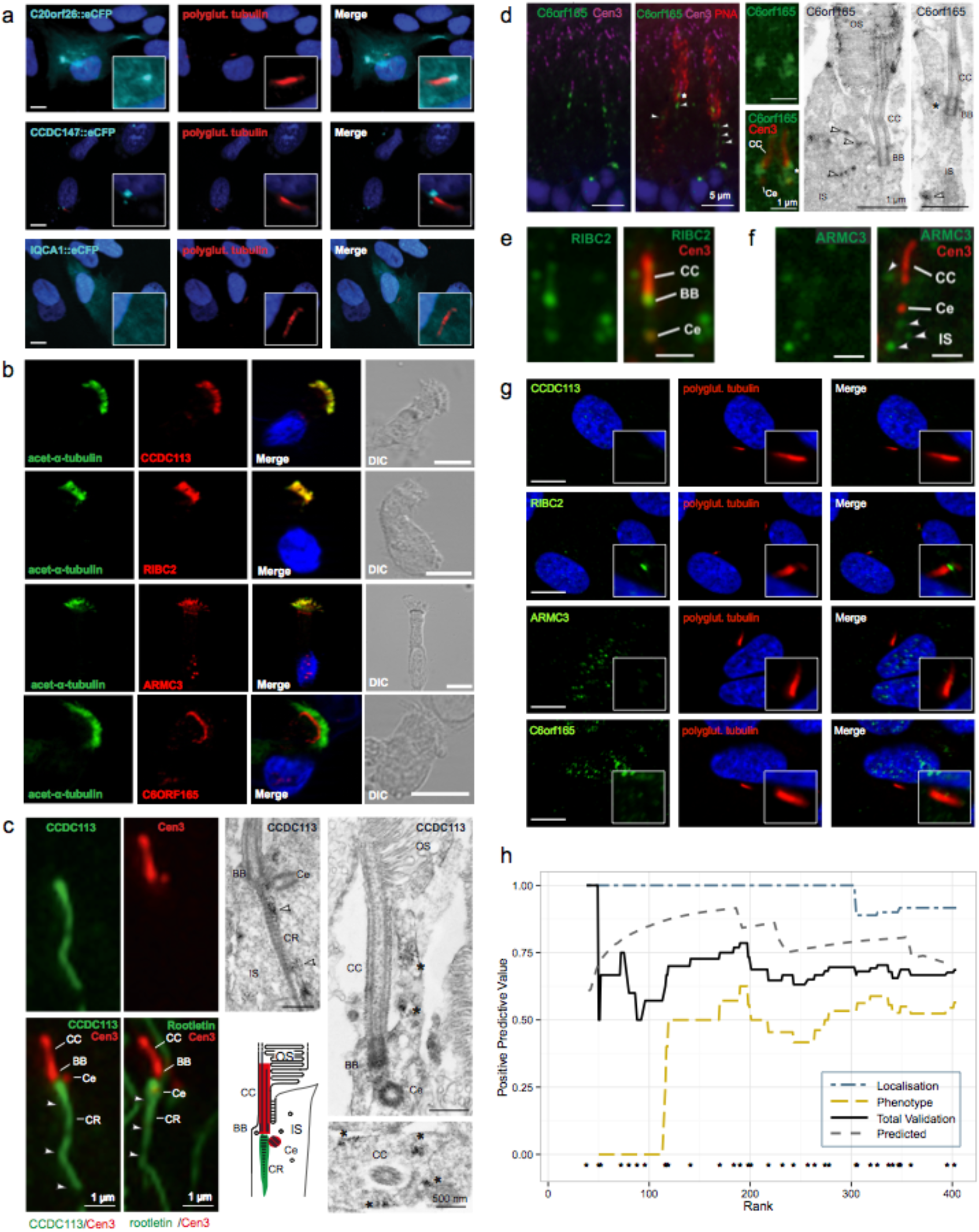
Validation by ciliary localization. a) Fluorescence microscopy of eCFP fused to C20orf26, CCDC147 and IQCA1 in hTERT-RPE1 cells. Acetylated alpha tubulin (red) is used to mark the axoneme. DAPI (blue) staining is used to mark the cell nucleus. IQCA1 does not appear to co-localize with acetylated alpha tubulin. b) Localization of CCDC113, RIBC2, ARMC3 and C6orf165 (red) compared with acetylated alpha tubulin (green) in human lung epithelial cells. c) Localization of CCDC113 in the primary sensory cilium of mature mouse photoreceptor cells. On the left: indirect 2-color immunofluorescence of CCDC113 (green) and centrin-3 (Cen3, red), a marker protein for the connecting cilium (CC), the basal body (BB) and the adjacent centriole (Ce), and of the ciliary rootlet (CR) marker rootletin (green) and Cen3 (red) indicates the localization of CCDC113 at the ciliary base and the CR projecting into the inner segment (IS). On the right: immunoelectron microscopy of CCDC113 confirms the localization of CCDC113 at the CR (arrowheads) and demonstrates accumulation of CCDC113 in the periciliary region of photoreceptor cells (asterisks). d) On the left: localization of C6orf165 in the primary sensory cilium of mature mouse cone photoreceptor cells. Indirect double immunofluorescence of C6orf165 (green) and Cen3 (magenta) in combination with the counterstaining with fluorescent PNA (red) revealed the localization of C6orf165 at the BB and the Ce of cone photoreceptors and a punctate staining in the IS (arrowheads). At the center: higher magnification of the double immunofluorescence of C6orf165 (green) and Cen3 (red). On the right: immunoelectron microscopy of the ciliary region of photoreceptors confirmed the ciliary and periciliary localization (asterisks) of C6orf165, but also demonstrated its presence in outer segments (OS) of cones. e) Double immunofluorescence of RIBC2 (green) and Cen3 (red) of a photoreceptor cilium showed localization of RIBC2 throughout the connecting cilium (CC) and the adjacent centriole (Ce) as seen by co-localization with the ciliary marker Cen3. f) Double immunofluorescence of ARMC3 (green) and Cen3 (red) of a photoreceptor cilium revealed the absence of ARMC3 from the CC (counterstained for Cen3) but a punctate staining in the periciliary region of the photoreceptor IS (arrowheads). g) Localization of CCDC113, RIBC2, ARM3 and C6orf165 compared with polyglutamylated tubulin in hTERT-RPE1 cells. h) Positive predictive value (PPV) of the Bayesian classifier based on the experimental validation outcomes plotted against CiliaCarta gene rank. The PPV of the combined validation converges to 0.67, which equals the predicted PPV (0.67, given 0.33 FDR). The asterisks (*) above the x-axis denote the ranks of the candidate genes and proteins tested for ciliary function or localization.

Four proteins (RIBC2, ARMC3, CCDC113 and C6orf165) were investigated for ciliary localization of the endogenous protein by immunofluorescence microscopy. RIBC2, ARMC3 and CCDC113 were observed to co-localize with acetylated alpha tubulin, a marker for the ciliary axoneme in human respiratory cells, while C6orf165 specifically localized to the base of the cilium (Fig. 4b). We also tested these four proteins in other model systems to investigate if their ciliary localization is model-specific. Location was validated in murine retinal sections by immunofluorescence and at the ultrastructural level via immunogold electron microscopy (Fig. 4c-f). All four proteins associated with the sensory cilia of photoreceptor cells. However, in hTERT-RPE1 cells only RIBC2 showed localization at the basal body suggesting that ciliary localization can be cell type specific (Fig. 4g). Concurrent to our efforts other labs recently identified ciliary involvement for RIBC2^26^ and CCDC113^27^.

#### Overall validation performance

Combining the results of all the validation experiments, we observed a ciliary localization or putative cilium-associated phenotype for 24 of the 36 candidates tested (Table 2). There appears to be a notable performance difference between the localization- and phenotype-based validation assays (PPV of 0.92 and 0.56 respectively, Fig. 4h). This difference is likely attributable to the fact that knockdown of a ciliary gene does not necessarily lead to an observable phenotype. Although we should keep in mind that the nematode and zebrafish ciliary phenotypes require confirmation via rescue experimentation, the overall experimentally determined positive predictive value (PPV) of the Bayesian classifier will be within 40 to 70%, approaching 67% depending on conservative or optimistic interpretations of the presence of false positives or false negatives in our validations(Fig. 4h).

The observed experimental PPV corresponds with the theoretical PPV of 67%, which corresponds to the FDR of 33% calculated for the candidate genes from the integrated CiliaCarta score (expected validation rate: 24.1 out of 36, p(x=24 given 33% FDR)=0.15, hypergeometric test, Fig. 4h). Even if we assume the lowest PPV of 40%, the experimental validation rate is significantly higher than expected by chance, based on the estimated prior distribution of ciliary versus non-ciliary genes in the human genome (expected 1.8 out of 36, p=3.34e-23 for PPV=67% and p=7.64e-10 for PPV=40%; hypergeometric test). The experimental verification of our candidate genes therefore validates our Bayesian classifier.

### OSCP1, a novel ciliary protein

Unbiased integration of large-scale genomics data can give rise to apparent inconsistencies with previous literature reports. For instance, organic solute carrier partner 1 (OSCP1, or oxidored-nitro domain-containing protein 1, NOR1), scores high on our CiliaCarta list (ranked 402), despite reports of varied functions not obviously consistent with a ciliary role, such as regulation of inflammation, apoptosis, proliferation and tumor suppression^28–30^. OSCP1 was first implicated as a tumor suppressor in nasopharyngeal cancer^28^ and was later found to modulate transport rates of organic solutes over the plasma membrane in rodents^31,32^. These two facets of OSCP1 function have never been connected to each other in literature. Subsequent independent research indicated a role for OSCP1 in regulation of inflammation and apoptosis^29^, and that it is specifically expressed in mouse testes^33^. The availability of a substantial set of literature on OSCP1 without any indication of ciliary involvement as predicted by our method, would have suggested that OSCP1 could have been a false positive in our Bayesian method. However, our *C. elegans* phenotype screen suggests that *C. elegans oscp-1* (R10F2.5) mutant alleles may possess modest sensory cilia defects (Fig. 2a & b). Therefore, we targeted OSCP1 for more detailed investigation of OSCP1 in *C. elegans*, zebrafish and mammalian cells.

### OSCP1 locates at the base and axoneme of the cilium in human, mouse and C. elegans cells

In *C. elegans*, expression of GFP-tagged OSCP-1, driven by its endogenous promoter, is restricted to ciliated sensory neurons (the only ciliated cells in the nematode), indicating a high likelihood of a cilium-associated function (Fig. 5a; Supplementary Fig. 5). Analysis of the subcellular localization pattern confirmed this by showing that OSCP-1::GFP localizes specifically to ciliary axonemes, including the ciliary base (Fig. 5a; Supplementary Fig. 5). In human hTERT-RPE1 cells, OSCP1::eCFP localizes at the basal body and daughter centriole in ciliated cells and at the centrioles of non-ciliated cells, indicating a basal body/centriole role for OSCP1 in this epithelial cell type (Fig. 5b). In some cells a clear punctate localization in the cytoplasm can also be observed.

**Figure 5.**
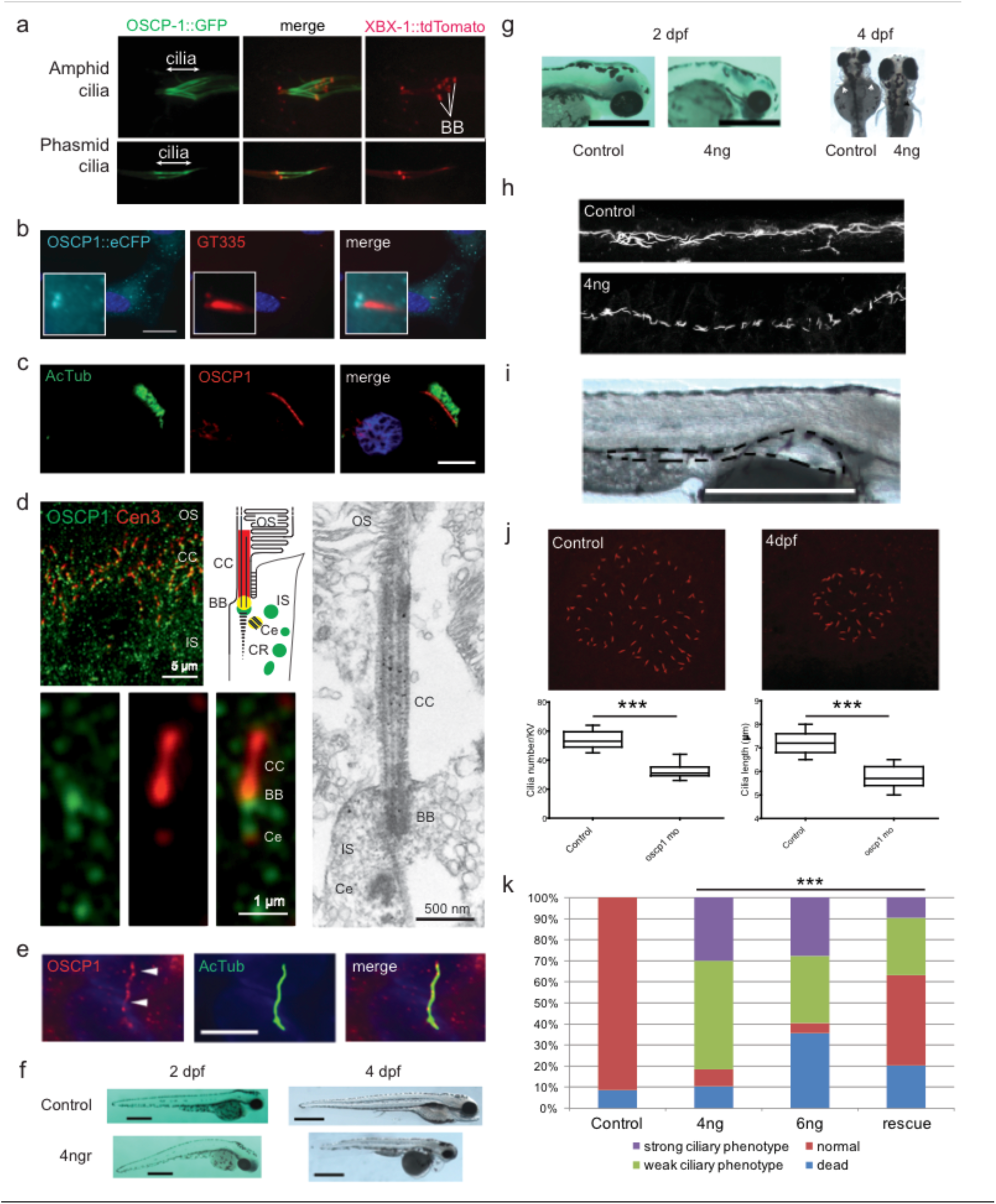
OSCP1 localizes to the cilium and regulates ciliary function in vivo. a) GFP-tagged OSCP-1 driven by its endogenous promoter is specifically expressed in ciliated sensory neurons in *C. elegans*, and the GFP-fusion protein is concentrated along the length of the cilium. Shown are fluorescent images of OSCP-1::GFP and XBX-1::tdTomato (ciliary marker) localisation in amphid and phasmid cilia. Basal bodies (bb) and cilia are indicated. b) OSCP1::eCFP localization in hTERT-RPE1 cells. OSCP1 localizes to the basal body and daughter centriole as well as in the cytosol in a punctate manner. c) OSCP1 localization in human respiratory cells (red) co-stained with acetylated tubulin (green). OSCP1 localizes to the cytosol, but specifically to the base of the ciliated crown of these multi-ciliated cells. d) Indirect high magnification immunofluorescence of OSCP1 (green) and centrin-3 (Cen3, red), a marker protein for the connecting cilium (CC), the basal body (BB) and the adjacent centriole (Ce), in the photoreceptor cilium region of an adult mouse. Immunoelectron microscopy of CCDC113 confirms the localization of OSCP1 at the base of the cilium. The schematic represents a zoom of the ciliary region of a photoreceptor stained for OSCP1 and Cen3 in the according colours. IS, inner segment; OS, outer segment. e) Immunostaining of serum starved murine IMCD3 cells with OSCP1 antibodies. OSCP1 is expressed in a punctate pattern along the axoneme. Acetylated tubulin and γ-tubulin (green), OSCP1 (Proteintech, 12598-1-AP, red) and nuclei (blue). Scale bar; 5 μm. f) Zebrafish embryos injected with 4 ng of oscpl splice morpholino 2 and 4 days post-fertilization (dpf). The characteristic ciliary phenotype with a curved body, small eyes and melanocyte migration defects at 2dpf. 4 dpf morphants display obvious pronephric cysts, small eyes, heart edemas, small heads and short bodies. Scale Bars 500 μm. g) Left panel: details of the head of 2 dpf zebrafish embryos showing small eyes and small head in the oscpl morphants. Scale bar 500 μm. Right panel: dorsal view of 4 dpf zebrafish embryos. Left embryo is a 4dpf oscpl morphant. Right embryo is a control. Scale bar 200 μm. Note the small eyes, melanocyte migration defects and small fin buds (white arrows) in the oscp1 morphants compared with the control fin buds (black arrows). h) Immunofluorescence staining of pronephric cilia at 24 hpf (acetylated a-tubulin). 4 ng oscpl morphants display shortened and disorganized cilia in the medial portion of the pronephric ducts. i) Detail of a pronephric cyst (outlined) in a 4 dpf zebrafish morphant. Scale bar 500 μm. j) Oscpl morphants Kupffer’s vesicle cilia staining. Oscpl morphants show smaller Kupffer’s vesicles with reduced cilia number per Kupffer’s vesicle (56 controls vs. 33 oscpl morphants) and shorter cilia (controls; 7.l μm vs oscpl morphants; 5.7 μm. Significance was determined by t-test p-value<0.0l. k) Dose dependent phenotype of oscpl morphants. After injecting 4 ng of oscpl morpholino the percentage of embryos with a weak phenotype is 5l% and embryos with a strong phenotype is 28%. Those percentages change when 6 ng of morpholino are used with 3l% of embryos showing with a weak phenotype and 27% with a strong phenotype, (strong and weak phenotypes described in Supplementary Fig. 7). The number of dead embryos increases when 6ng of oscpl morpholino are used (38%) compared with 4 ng of oscpl morpholino (l0%). We injected zebrafish embryos at one cell stage with 4 ng of oscpl morpholino and l00 pg of human OSCPl mRNA. The rescue increased the normal phenotype percentage from 8% to 43% and decreased the weak phenotype from 5l% to 27% and the strong phenotype from 28% to 9.5%. Significance was determined by χ2 test, p<0.000l.l

To assess the localization of the endogenous OSCP1 protein in various mammalian cells, we used commercially available antibodies from three suppliers (Proteintech, Atlas and Biorbyt). In human multi-ciliated respiratory cells, OSCP1 shows an increased concentration at the base of the cilia (Fig. 5c). In murine photoreceptors OSCP1 is localized at the inner segments (Fig. 5d), where it is particularly abundant at the base of the connecting cilium (the equivalent of the transition zone in primary cilia), and at low concentration at the adjacent centriole. Immunoelectron microscopy analysis also shows OSCP1 at the basal body, occasionally at the connecting cilium and sporadically within the inner segment (Fig. 5d). Serum-starved IMCD3 cells (murine collecting duct cells) showed ubiquitous punctate cytoplasmic staining with increased signal at the basal body and along the ciliary axoneme (Fig. 5e; Supplementary Fig. 6). In ATDC5 cells (murine pre-chondrocyte cells) OSCP1 is mainly localized to the ciliary axonemes, although the ciliary base signal is less apparent (Supplementary Fig. 6).

Together, these exhaustive subcellular localization analyses place OSCP1 at ciliary structures, including the ciliary base and axoneme, although there are some subtle differences between cell types. We also frequently observed extensive non-ciliary signals for OSCP1, consistent with published reports for OSCP1 localizations at other organelles (ER, Golgi, Mitochondria) and within the cytosol^34,35^.

### OSCP1 is required for cilium formation in multiple zebrafish tissues, but dispensable in C. elegans sensory neurons

To investigate possible ciliogenesis roles for OSCP1, we first examined cilia in zebrafish morphants depleted for *oscp1* in fertilized eggs. In embryos at 2 days post fertilization (dpf) developmental phenotypes were observed that are often associated with ciliary defects: a curved body axis, small eyes and melanocyte migration defects (Fig. 5f). Likewise, the 4 dpf morphants presented with pronephric cysts, small eyes, heart edema, small heads and short bodies (Fig. 5f & g). The cilia in the medial portion of the pronephric ducts of *oscp1* morphants were shortened and disorganized (Fig. 5h&i), and in the Kupffer's vesicle cilium length and number was decreased compared to control injected larvae (Fig. 5j). Co-injecting human *OSCP1* mRNA together with the morpholino partially rescued the observed phenotype, indicating that the observed phenotype is specific for loss of *oscp1*function (Fig. 5k and Supplementary Fig. 7). Therefore, in zebrafish, *oscp1* is required for cilium formation and associated functions in many tissue types and organs.

In contrast, analysis of the *oscp-1*(*gk699*) null allele in *C. elegans* revealed that OSCP-1 is not required for cilium formation. Using fluorescence reporters and transmission electron microscopy, the amphid and phasmid channel cilia appeared to be intact, and full length (Supplementary Fig. 5 and 9). In addition, *oscp-1* does not appear to be functionally associated with the transition zone at the ciliary base; disruption of this prominent ciliary domain (‘gate’) in the *mks-5* mutant^36^ does not affect OSCP-1::GFP localization, and loss of *oscp-1* itself does not influence the localization of several transition zone proteins (Supplementary Fig. 5). Thus, OSCP1 is differentially required for cilium formation in worms and zebrafish, reflecting species distinctions in the ciliary requirement for this protein. These distinctions could reflect redundancy of OSCP1 function with another ciliary protein in the nematode, but not in zebrafish, or differences in cell type requirements (nematode sensory neurons versus zebrafish epithelial cells). Clearly, based on its restricted expression in ciliated cells and localization with the ciliary axoneme, *C. elegans* OSCP-1 is serving a ciliary function, although the specifics of this role remain to be elucidated.

## Discussion

With the current interest in cilia biology it is certain that many new genes will be implicated in ciliary function for several years to come. With the advent of systems biology and the need to understand the cilium as a whole the research community requires an inventory of genes and proteins involved in ciliary structure and function. The obvious sources for such an inventory, GO^4^ and the SCGS^5^, currently only cover 608 human genes (510 in GO as of December 3rd 2015, 302 for the SCGS). By applying a naive Bayesian integration of heterogeneous large-scale ciliary data sets we have expanded this set by 38% to 836 human genes, adding 228 putative genes to the known cilium gene repertoire (Supplementary Fig. 8). GO term enrichment analysis indicates that these putative ciliary genes are enriched for genes that are “unclassified” (86 genes, 1.69 fold enrichment, p-value < e-100) suggesting possible new ciliary biology to be discovered. We put forward these putative ciliary genes, together with the SCGS and the GO annotated genes, as the “CiliaCarta”, a compendium of ciliary components with an estimated FDR of 10% (Supplementary table 3). This community resource can be used to facilitate the discovery of new cilium biology and to identify the genetic causes of cilia related genetic disease. CiliaCarta therewith has a very different purpose than the SCGS, it serves as a tool for discovery of new genes, rather than as a reference of known cilium genes.

CiliaCarta is still likely to be incomplete. Estimates of the ciliary proteome range from one to two thousand proteins depending on the techniques used and on the types of cilia and the species studied^19,37^. In addition to obtaining the CiliaCarta list of proteins, our Bayesian analysis allows us to obtain an objective estimate of the total number of ciliary proteins. Using the posterior probabilities and the outcome from the validation experiments we estimate the size of the ciliome to be approximately 1200 genes (Methods).

Predicting the genes responsible for an organelle structure or function poses the question of where we draw the boundary between that organelle and the rest of the cell. For example, does the basal body in its entirety belong to the cilium or are some components to be considered exclusive to the centrioles? Furthermore, one can argue that in the case of the cilium, proteins that are not part of the cilium, but play a role in the transport of proteins to the ciliary base or regulation of cilium gene expression, can be regarded as components of the ‘ciliary system’. In practice, what we regard here as a ciliary component depends on the definition within the SCGS, which is used to weigh the data sets, and on our experimental validations. In both we have taken a rather inclusive approach by regarding genes whose disruption cause ciliary phenotypes as ciliary genes. This approach makes our predictions relevant to human disease. Indeed KIAA0753, which falls within our 25% cFDR list of cilium genes, was recently shown to interact with OFD1 and FOR20 at pericentriolar satellites and centrosomes, and gives rise to oral-facial-digital syndrome^38^, a phenotype associated with disruption in the ciliary transition zone and the basal body.

Data integration through objective quantification of the predictive value of individual large-scale data sets allows one to find new functions and associations, without bias from previous studies. As a case in point we report here that OSCP1, not previously implicated in ciliary functions based on the existing published literature, is validated as a ciliary protein in four species as determined by multiple independent experimental methods. Re-evaluation of previous experimental evidence on OSCP1 function does not exclude ciliary involvement. Nevertheless, the cilium contains several specific ion-channels in its membrane^39^ and the organelle has been implicated to play a role in the development of cancer^40–42^. Therefore connecting OSCP1 to the cilium might provide the missing link to connect previously observed effects of OSCP1 on organic solute in-/efflux and its role in nasopharyngeal cancer. Our results therefore provide a cellular target and biomolecular framework to further unravel OSCP1 function.

Systematic integration of heterogeneous large-scale cilia data sets by employing Bayesian statistics combined with medium throughput experimental validation is a powerful approach to identify many new ciliary genes and provides a molecular definition of the cilium. The experimental observations of a potential ciliary role for selected high-confidence candidates, together with the results from the cross validation, indicates that the top tier of the entire ranked human genome is highly enriched for ciliary genes. The genome-wide CiliaCarta Score and ranking as provided here, should therefore make it possible to efficiently and objectively prioritize candidate genes in order to discover new ciliary genes and ciliary functions.

## Materials and methods

### Data set collection and mapping

All data sets were mapped to the ENSEMBL human gene set version 71, release April 2013^43^. Resources using other identifiers (i.e. Entrez, Uniprot) were mapped to ENSEMBL gene IDs using ENSEMBL BioMART (version 71). Orthologs from non-human data sets were mapped using the ENSEMBL Compara ortholog catalogue from ENSEMBL 71, with exception of Sanger sequences from Y2H screens based on the Bovine cDNA library that were mapped to Bovine genome sequences by the BLAT tool^44^ in the UCSC genome browser^45^ and subsequently mapped to human orthologs using the ‘non-cow RefSeq genes track’ from this browser.

### DNA constructs

Bait protein selection was based on the association of proteins with ciliopathies (including mutant vertebrates showing ciliopathy features), involvement in IFT or part of our candidate list of ciliary proteins. Gateway-adapted cDNA constructs were obtained from the Ultimate™ ORF clone collection (Thermo Fisher Scientific) or generated by PCR from IMAGE clones (Source BioScience) or human marathon-ready cDNA (Clontech) as template and cloning using the Gateway cloning system (Thermo Fisher Scientific) according to the manufacturer’s procedures followed by sequence verification.

### Yeast two-hybrid system

A GAL4-based yeast two-hybrid system was used to screen for binary protein-protein interactions with proteins expressed from several different cDNA libraries (see below) as described previously^46^. Yeast two-hybrid constructs were generated according to the manufacturer’s instructions using the Gateway cloning technology (Thermo Fisher Scientific) by LR recombination of GAL4-BD Gateway destination vectors with sequence verified Gateway entry vectors containing the cDNA’s of selected bait proteins.

Constructs encoding full-length or fragments of bait proteins fused to a DNA-binding domain (GAL4-BD) were used as baits to screen human oligo-dT primed retinal, brain (Human Foetal Brain Poly A+ RNA, Clontech), kidney (Human Adult Kidney Poly A+ RNA, Clontech) or testis cDNA libraries, or a bovine random primed retinal cDNA library, fused to a GAL4 activation domain (GAL4-AD). The retina and testis two-hybrid libraries were constructed using HybriZAP-2.1 (Stratagene), the brain and kidney two-hybrid libraries were constructed using the “Make Your Own Mate & Plate™ Library System” (Clontech).

The yeast strains PJ96-4A and PJ96-4α (opposing mating types), which carry the *HIS3* (histidine), *ADE2* (adenine), *MEL1* (α-galactosidase), and *LacZ* (β-galactosidase) reporter genes, were used as hosts. Interactions were identified by reporter gene activation based on growth on selective media (*HIS3* and *ADE2* reporter genes), α-galactosidase colorimetric plate assays (*MEL1* reporter gene), and β-galactosidase colorimetric filter lift assays (*LacZ* reporter gene).

### Affinity purification of protein complexes using SILAC

#### DNA constructs and cell culture

Experiments were essentially performed as described before^47^. In short, N-terminally SF-TAP-tagged bait proteins that were obtained by LR recombination of TAP-destination vector with sequence verified Gateway entry vectors containing the cDNA’s of selected bait proteins using Gateway cloning technology (Thermo Fisher Scientific). HEK293T cells were seeded, grown overnight, and then transfected with SF-TAP-tagged bait protein constructs using Effectene (Qiagen) according to the manufacturer’s instructions. HEK293T cells were grown in SILAC cell culture medium as described^47^.

#### Affinity purification of protein complexes

For one-step Strep purifications, SF-TAP-tagged proteins and associated protein complexes were purified essentially as described previously^47^. In short, SILAC labeled HEK293T cells, transiently expressing the SF-TAP tagged constructs were lysed in lysis buffer containing 0.5% Nonidet-P40, protease inhibitor cocktail (Roche), and phosphatase inhibitor cocktails II and III (Sigma-Aldrich) in TBS (30 mM Tris-HCl, pH 7.4, and 150 mM NaCl) for 20 minutes at 4°C. After sedimentation of nuclei at 10,000 g for 10 minutes, the protein concentration was determined using a standard Bradford assay. Equal protein amounts were used as input for the experiments to be compared. The lysates were then transferred to Strep-Tactin-Superflow beads (IBA) and incubated for 1 hour before the resin was washed 3 times with wash buffer (TBS containing 0.1% NP-40 and phosphatase inhibitor cocktails II and III). The protein complexes were eluted by incubation for 10 minutes in Strep-elution buffer (IBA). The eluted samples were combined and concentrated using 10-kDa cutoff VivaSpin 500 centrifugal devices (Sartorius Stedim Biotech) and prefractionated using SDS-PAGE and in-gel tryptic cleavage as described elsewhere^48^.

#### Quantitative mass spectrometry

After precipitation of the proteins by methanol-chloroform, a tryptic in-solution digestion was performed as described previously^49^. LC-MS/MS analysis was performed on a NanoRSLC3000 HPLC system (Dionex) coupled to a LTQ OrbitrapXL, respectively coupled to a LTQ Orbitrap Velos mass spectrometer (Thermo Fisher Scientific) by a nano spray ion source. Tryptic peptide mixtures were automatically injected and loaded at a flow rate of 6 μl/min in 98% buffer C (0.1% trifluoroacetic acid in HPLC-grade water) and 2% buffer B (80% acetonitrile and 0.08% formic acid in HPLC-grade water) onto a nanotrap column (75 μm i.d. × 2 cm, packed with Acclaim PepMap100 C18, 3 μm, 100 Å Dionex). After 5 minutes, peptides were eluted and separated on the analytical column (75 μm i.d. × 25 cm, Acclaim PepMap RSLC C18, 2μm, 100 Å Dionex) by a linear gradient from 2% to 35% of buffer B in buffer A (2% acetonitrile and 0.1% formic acid in HPLC-grade water) at a flow rate of 300 nl/min over 33 minutes for EPASIS samples, and over 80 minutes for SF-TAP samples. Remaining peptides were eluted by a short gradient from 35% to 95% buffer B in 5 minutes. The eluted peptides were analyzed by using a LTQ Orbitrap XL, or a LTQ OrbitrapVelos mass spectrometer. From the high-resolution mass spectrometry pre-scan with a mass range of 300-1,500, the 10 most intense peptide ions were selected for fragment analysis in the linear ion trap if they exceeded an intensity of at least 200 counts and if they were at least doubly charged. The normalized collision energy for collision-induced dissociation was set to a value of 35, and the resulting fragments were detected with normal resolution in the linear ion trap. The lock mass option was activated and set to a background signal with a mass of 445.12002^50^. Every ion selected for fragmentation was excluded for 20 seconds by dynamic exclusion.

For quantitative analysis, MS raw data were processed using the MaxQuant software^51^ (version 1.5.0.3). Trypsin/P was set as cleaving enzyme. Cysteine carbamidomethylation was selected as fixed modification and both methionine oxidation and protein acetylation were allowed as variable modifications. Two missed cleavages per peptide were allowed. The peptide and protein false discovery rates were set to 1%. The initial mass tolerance for precursor ions was set to 6 ppm and the first search option was enabled with 10 ppm precursor mass tolerance. The fragment ion mass tolerance was set to 0.5 Da. The human subset of the human proteome reference set provided by SwissProt (Release 2012_01 534,242 entries) was used for peptide and protein identification. Contaminants like keratins were automatically detected by enabling the MaxQuant contaminant database search. A minimum number of 2 unique peptides with a minimum length of 7 amino acids needed to be detected to perform protein quantification. Only unique peptides were selected for quantification.

### Protein-protein interaction data processing

Based on three ciliary protein-protein interaction (PPI) data sets, we inferred proteins to “interact with ciliary components” as a proxy for being part of the cilium. First, we obtained protein complex purification data from a large-scale study on the identification of ciliary protein complexes by tandem-affinity purification coupled to mass spectrometry (TAP-MS) for 181 proteins known or predicted to be involved in ciliary functions^17^. Since integration of this data set into the CiliaCarta many pull-downs were repeated and reverse experiments included. As a result, our data set includes 539 found interactors that are not part of the now published final data set (4702 proteins, Table S2 from Boldt *et al*.^17^), and does not include 679 new interactors that have been identified since June 2013. The complete and current data is available at http://landscape.syscilia.org/ and IntAct [IM-25054]. Second, we obtained affinity purification data for 16 bait proteins from a more sensitive and quantitative approach using affinity purification combined with stable isotope labeling of amino acids in cell culture (SILAC). In total 1301 interactors were identified by SILAC in 57 experiments (Supplementary table 4). Third, we also obtained direct protein-protein interaction data from several independent yeast two-hybrid (Y2H) screens against cDNA libraries derived from hTERT-RPE1 (retinal pigment epithelial) cells, as well as brain, kidney, retina and testis tissue. In total 69 Y2H screens were performed using 27 baits, identifying a total of 343 interacting proteins (Supplementary table 5).

The SILAC and Y2H studies were focused on finding new interactors for selected ciliary proteins of interest and were not part of a systematic analysis. Parts of the resulting PPIs were published in previous studies (four out of 16 baits for SILAC^47,52,53^ and nine out of 27 baits for Y2H^53-61^), however here we consider the entire PPI data sets. The complete data sets are publicly available in Supplementary tables 4 & 5 and at the IntAct database^62^ under references {DB reference 1} (SILAC) and {DB reference 2} (Y2H) (note to reviewers: datasets will be submitted to IntAct before publication). Because the TAP-MS and SILAC data sets are based on similar methodology and have a large bait overlap (14 out of 16 SILAC baits were used in the TAP-MS data set), we merged them into a single data set (Mass-spec based PPI) with 4799 unique proteins identified to interact with 184 bait proteins.

The Y2H, SILAC and TAP-MS data were transformed to genome wide data sets by defining genes as (“found”) when the gene product was found to interact with the baits, and as 0 (“not found”) when the gene product was not found to interact. Due to the large overlap in baits and the largely similar methods used in the TAP-MS and SILAC data sets we decided to combine the data sets in order to avoid counting the interacting proteins multiple times and thereby artificially overestimating their CiliaCarta Scores. The mass-spectrometry data sets and Y2H data sets were found to be sufficiently different to include them as separate data sets (positive set correlation is 0.13, Supplementary Fig. 10).

### Expression screen data set

In expression screening^63^ separate gene-expression data sets are weighted for their potential to predict new genes for a system by measuring, per data set, the level of co-expression of the known genes. We have already successfully applied this method to predict TMEM107 as part of the ciliary transition zone ^64^ and now extend the approach to the complete cilium. An integrated cilium co-expression data set was constructed by applying the weighted co-expression method WeGet^65^ to ciliary genes in 465 human expression data sets available in the NCBI Gene Expression Omnibus^66^. For individual genes, correlations of their expression profiles were determined with expression profiles of the set of ciliary components. The contribution of each data set to a final co-expression score per gene was weighed by how consistently the set of cilia components were expressed together, i.e. how well the data set in question is able to detect ciliary components^65^. To avoid circularity with the training of the Bayesian classifier the expression screen was performed using a gene set of ciliary components from GO (GO:Cilium) and removed any overlap from the positive set used to evaluate this data set for the Bayesian classifier. The data set from Ross *et al.^20^*,which has been included in the Bayesian classifier, has been excluded from the microarray data sets used in the expression screen.

### Ciliary co-evolution data set

Given the large number of independent losses of the cilium in eukaryotic evolution (we counted eight independent loss events throughout the eukaryotic kingdom)^67–69^, presence/absence profiles have a high value for predicting new cilium genes^12,64^. We constructed a comprehensive co-evolution data set from a comparative genomics analysis of presence-absence correlation patterns over a representative data set of eukaryotic species. We correlated the occurrence of orthologs of 22,000 human genes in 52 eukaryotic genomes to that of cilia or flagella using differential Dollo parsimony (DDP)^70^. A perfectly matched profile pair would obtain a DDP of 0 (all events match, no differences), while mismatching profile pairs would receive a DDP equal to the number of evolutionary events that did not occur at the same time in evolution (e.g. the gene was lost in a lineage still maintaining a cilium, or the gene was maintained in a lineage in which the cilium has been lost). Thus we obtained an objective measure for each human gene that describes how well its evolutionary trajectory (i.e. point of origin and independent loss events) matches that of the ciliary system. A number of mismatches are expected for some ciliary genes; *Plasmodium falciparum* for instance has maintained a cilium, but has lost all genes of the IFT machinery. Due to the topology of the species tree we observed a complicated distribution of genes from the positive and negative sets (Supplementary Fig. 11a): for low DDP scores (0-6) we did not observe a single negative gene, which would result in unrealistic log odd scores (i.e. infinity). To avoid these unrealistic log odds, we decided to combine the DDP scores into two categories, namely genes with a DDP ≤ 9 and genes with a DDP ≥ 10. Genes with a score between 0 and 9 were generally overrepresented among ciliary genes (Supplementary Fig. 11b).

### Transcription factor binding sites data set

The RFX and FOXJ1 transcription factors play an important role in the regulation of ciliogenesis^71,72^. X-box (RFX) or a FOXJ1 transcription factor binding site (TFBS) have been used to predict novel ciliary genes in *Caenorhabditis elegans* (nematode) and *Drosophila melanogaster* (fruit fly)^73–75^. We processed the publicly available data sets from the 29 mammals project^18^ to obtain human genes with a conserved X-box or FOXJ1 TFBS in their promoters, which were defined as 4 kilobase (kb) windows centered (i.e. 2kb upstream and 2kb downstream) at all annotated transcription start sites of the gene. The restriction that the TFBS motifs are conserved among mammalian species infers a higher level of confidence that these motifs are indeed relevant and not spurious hits. The final data set was constructed by defining two categories, namely: “Gene has a X-box and/or FOXJ1 TFBS”, represented as 1, and “Gene does not have a X-box and/or FOXJ1 TFBS”, represented as 0. We found relatively limited overlap with the previous invertebrate X-box TFBS data sets: 13% of the genes with an X-box in *C. elegans* (225 out of 1695 genes)^73–75^ and 15% in *D. melanogaster* (71 out of 470)^75^ have a conserved X-box in human. This low overlap may result from differences in the sensitivity detecting functional X-box sequences. It might also indicate that the X-box motifs are transient in the genome; i.e. often gained and lost, as has also been observed in vertebrate evolution of other TF binding sites^76^.

### Published data sets

There are a number of high-throughput cilia data sets available from the CilDB ^15^ that can complement the data sets mentioned above. We only included data sets from mammalian species to avoid significant issues with orthology (i.e. avoid mapping to paralogous genes) and which had a broad coverage of the entire genome/proteome (Supplementary fig. 1). We excluded data sets that focused specifically on the centriole/basal body, since this structure is also affected by the cell cycle^77,78^ and therefore could potentially skew the Bayesian classifier towards this process. We avoided redundancy in the data sets by selecting only one data set per experiment type (e.g. proteomics, expression). We only considered proteomics datasets specifically generated for the cilium as opposed to whole cell proteomics data sets to minimize false positives. The data from Ross *et al.^20^* (expression) and Liu *et al.^19^* (proteomics) were selected based on these criteria. These data sets were extracted from CilDB^15^ and implemented using the predefined confidence categories from CilDB (Low confidence, medium confidence, high confidence). Genes not covered by these data sets were assigned to the “not found” category.

### Training sets

We used the SYSCILIA Gold Standard (SCGS)^5^ as our positive training set. We did not use GO annotations as we regard the SCGS, that has been annotated by experts in the cilium field, of higher quality Furthermore, having an independent cilium genes dataset allowed us to prevent circularity in the expression screening analysis (see below). We are currently in the process of improving cilia-related GO terminology and transferring our SCGS annotations^79^. The SCGS, or positive set, contains a total of 302 manually curated human ciliary genes. We constructed a negative set by selecting genes annotated in GO to function in processes and cellular compartments we deemed least likely to be (in)directly involved in ciliary processes. We selected genes annotated with at least one of the following GO Cellular Component terms: extracellular, lysosome, endosome, peroxisome, ribosome, and nucleolus. We ensured that the positive and negative training sets do not overlap by removing genes found in both from the negative set. Since the majority of human genes are expected to be non-ciliary the similarity between the score distribution of the negative set and the remaining “other genes” indicates that the negative set overall gives an excellent representation of what we reasonably can expect to be non-ciliary genes. The final negative set contains 1275 genes.

We adapted the positive training set for the PPI data sets as well as the expression screen data set to avoid overtraining. Since many of the baits used in the PPI data are known ciliary components and therefore part of the positive training set, this could lead to a potential overestimation of the predictive value of the data sets. Therefore we excluded the bait proteins from the positive set for evaluating the Y2H, TAP-MS and SILAC data. The training of the expression screen was performed using a gene set of ciliary components from GO (GO:Cilium) and to avoid overtraining we subtracted the overlap from the positive set used for the Bayesian classifier. In this way we avoided inflation of the predictive values for these data sets.

### Bayesian classifier

#### Performance and false discovery rate calculations

The performance of each data set for predicting ciliary genes as well as the integrated Bayesian classifier was evaluated using the positive and negative training sets. We determined the fraction, or recall, of the positive training set, which is also known as the sensitivity or true positive rate (TPR).

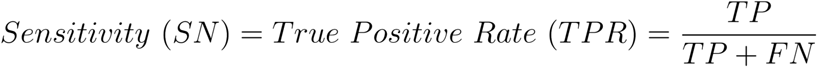
Where True Positives (TP) are true ciliary genes that are correctly discovered by the data set, and False Negatives (FN) are true ciliary genes that were not discovered by the data set. We also determined the fraction of the negative set retrieved by the data set, also known as the false positive rate (FPR).

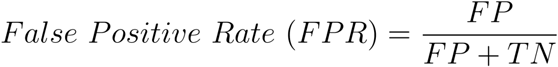
Where True Negatives (TN) are non-ciliary genes that are correctly excluded from the data set, and False Positives (FP) are non-ciliary genes incorrectly discovered in the data set. The FPR together with the TPR are used to calculate the predictive ability for each data set (see below).

The FPR is related to the specificity (SP), another well-known metric for the quality of a data set, as follows:

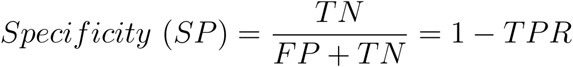
The false discovery rate (FDR) denotes the chance of encountering a false positive among a set of predictions and thus reflects the trustworthiness of a prediction. The FDR is given by:

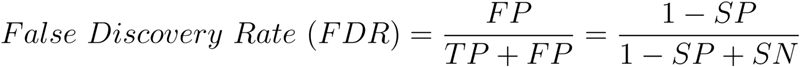
We use the FDR to determine the overall performance of the classifier given a specified threshold of the CiliaCarta Score. However, the FDR depends on both training sets and is sensitive to deviations in set size compared to the actual populations of positives and negatives for the whole genome, and thus we need to correct the canonical FDR equation for the differences in the population and set size:

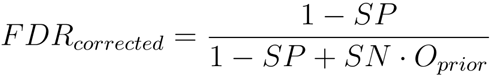
Throughout the text we refer to this adjusted FDR as the corrected FDR (cFDR). The positive predictive value (PPV) is directly related to the FDR and reflects the probability that a gene within the set threshold will indeed be ciliary:

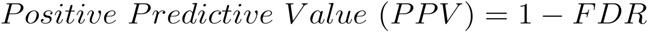
The PPV and FDR therefore reflect the chance of success or failure within a set of genes rather than for individual genes; i.e. they do not reflect the per gene probability for it being ciliary or not. Instead, the per-gene probabilities are reflected by the posterior odds, the CiliaCarta Score, see below.

At a cFDR cut-off of 25% our rank-ordered list of predictions covers 404 genes. However, this set contains genes from both the negative and the positive training sets, and after excluding those we obtain 285 predictions. This remaining set will however have a different FDR since we removed the training set genes. We obtain the FDR for the remaining 285 genes by: (i) calculating the number of TP and FP based on the cFDR threshold, (ii) subtracting the number of genes from the training sets (TP - genes in positive set, FP - genes in negative set) and (iii) recalculating the FDR using these adjusted TP and FP values. This results in a FDR of 33% for the 285 candidate predictions.

In our experimental validations 24 out of 36 genes are positive for ciliary phenotype and/or localization. We can derive an observed FDR directly from these numbers, i.e. 24 TP and 12 FP. The observed FDR is thus 12/36 = 33%, the same as the estimated FDR for our candidate genes.

#### Calculation of the CiliaCarta Score using naive Bayesian integration

Naive Bayesian integration allows a direct comparison and weighing of many and diverse data sets describing the properties of ciliary genes and integrates these into a single probabilistic score for each gene accommodating for missing data^7,8,80,81^. This approach has been successfully used to predict for instance mitochondrial^81^ and innate immunity^8^ genes.

For a given gene in the human genome we calculate the conditional probability that the gene is involved in ciliary processes given the observed evidence in the data sets. For our purposes, since we have only two possible outcomes (i.e. ciliary vs. non-ciliary) it is more convenient and appropriate to use odds instead. We can write the probability that a gene is ciliary given the outcome of the experiment in data set 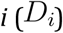 for all data sets j:

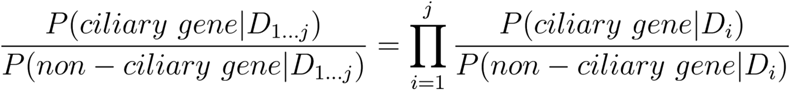
These odds cannot be calculated directly, but we can obtain them using Bayes’ theorem by approximating the reverse likelihood ratio *L* that a gene is observed in the data sets given it is either ciliary, or non-ciliary:

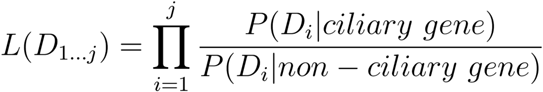
We can calculate this likelihood directly from the distribution of the training sets. This equates to the ratio between the data sets’ true positive rate for ciliary genes (i.e. the proportion of known ciliary genes retrieved), and the false positive rate for non-ciliary genes (i.e. the proportion of known non-ciliary genes retrieved). Using Bayes’ theorem we can now obtain the final (posterior) odds from the likelihood ratio *L* as follows:

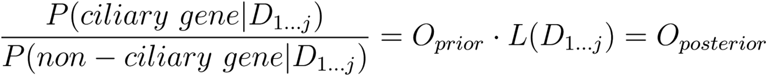

Where 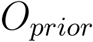 is the prior odd, i.e. the odds for a gene to be ciliary if one would randomly sample from the genome:

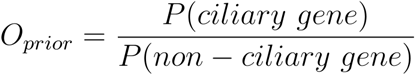
And thus we obtain the posterior probability for data sets *D_1-j_*:

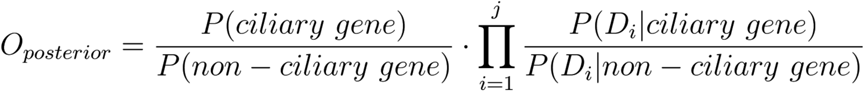

Finally we obtain the CiliaCarta Score by log_2_ transformation of the individual terms to get an additive score:

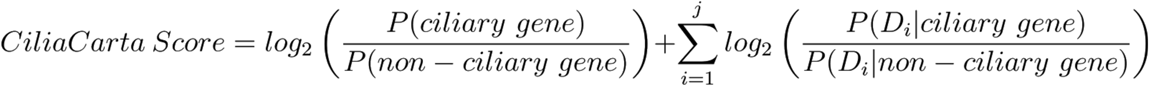
The additive nature of the log transformed posterior odds equation makes the contribution of each data set to the final score more insightful. It also makes the score robust against rounding errors otherwise encountered during multiplication of the untransformed odds. The log-odds for each individual data set per sub-category are listed in Supplementary table 7.

#### Determining the prior

One of the elements in Bayesian calculations is estimating the prior: the *a priori* expectation of how many ciliary genes there are in the human genome. Although the ranking of genes does not depend on the prior, it is required to obtain a corrected false discovery rate (see above). Furthermore it gives a meaning to the posterior log odd scores (the CiliaCarta score), i.e. a positive log odd means that the gene is more likely to be ciliary than non-ciliary and a negative log odd means that it is more likely to be non-ciliary.

To our knowledge there is no substantiated estimate for the number of genes involved in the cilium. Currently 608 human proteins have been annotated as being part of the cilium in a combination of GO (G0:0005929 Cell Component Cilium & G0:0042384 Biological Process Cilium Assembly, Ensembl biomart as of December 3rd 2015) and the SCGS, but we can reasonably assume the total number of ciliary genes to be much higher. For instance, the ciliary proteome as identified by Liu *et al*. in the mouse photoreceptor sensory cilium entails 1185 to 1968 proteins, depending on the stringency of the filters applied^19^. We have chosen a prior that we deem to be both reasonable and conservative: 5% of the human genome (i.e. 1135 ciliary genes). The 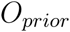 then becomes:

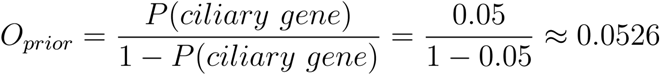

#### Conditional independence

An assumption of our naïve Bayesian approach is that the outcome of one data set is independent of the outcome of another. This assumption of independence is not always attainable, since for instance gene expression is to some extent biologically correlated to the presence of proteins in the proteomics data sets. Violations of the independence assumption can bias the predictions and can lead to an overestimation of the likelihood scores. However previous work has shown that, regardless of biological correlations between data sets, naive Bayesian integration of genomics data is highly effective to predict novel genes involved in a molecular system^7,81^. Analysis of the correlations suggests that the data sets used to predict ciliary genes are largely complementary (Supplementary Fig. 10). Several data sets have high correlations, such as ciliary co-evolution and co-expression. However these data sets are methodologically and experimentally completely unrelated and thus the high correlation is purely based on the ability of the methods to predict ciliary genes.

#### Ciliome size estimation

The Bayesian framework can be used to obtain a systematic estimate for the total number of ciliary genes in two ways. The first approach involves fitting a new 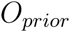 based on the observed validation rate of our experiments; that is, we make use of the discrepancy between the expected and experimentally determined number of hits. We can estimate this new 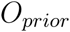 by equating the c FDR of the Bayesian integration, which depends on our original 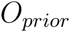 to the observed FDR of the validation experiments:

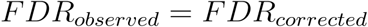

Where 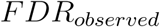 is calculated from the TP and FP determined from the validation experiments (24 and 12 resp.). The 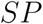 and 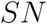 for the cFDR can be obtained from the training sets. We then try to find an 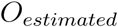 for which the following equation holds:

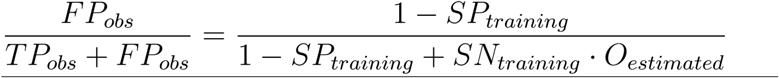
Atypically, in our study the observed FDR and the cFDR are equal (i.e. 33%), which indicates that 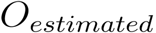 essentially equals our initially chosen 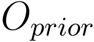 (1135 ciliary genes).

The second approach to estimate the ciliome size is based on the Bayesian posterior probabilities obtained for each gene (the CiliaCarta Scores). To determine the expected value (i.e. the number of positives) among a set of ciliary candidates, we considered each gene to be a random variable with a binary outcome (success, a true ciliary gene, or failure, not a ciliary gene) whose probability of success is defined by 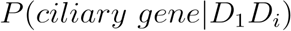. Effectively this corresponds to a Bernoulli process with different probabilities for success or failure for each successive trial. Assuming independence, the expected value *E* for a set of *n* binary random variables (i.e. ciliary candidates) is equal to the sum of the expected values of the individual variables. The expected value for an individual binary random variable, in turn, equals its probability of success. From the posterior odds:

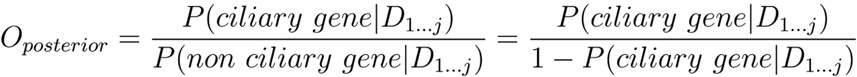
We can obtain the probability that a gene is ciliary by rewriting the above as:

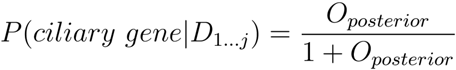
Since the CiliaCarta Score (CCS) is the log_2_ of the posterior odds we finally get

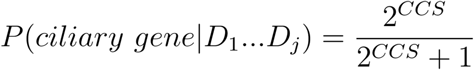
And thus to obtain the expected value E becomes:

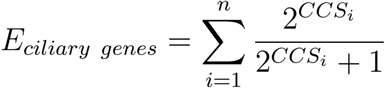

The expected value for the number of ciliary genes based on the Bayesian posterior probabilities was found to be 1273. It should be noted that the posterior CiliaCarta probabilities, and hence the expected value for a set of genes, depend on the prior estimation of the number of ciliary genes. We have above already established that our chosen prior was accurate based on our validation outcome. Indeed the posterior expected number of ciliary genes is close to the prior expected number of ciliary genes (1135, difference of 138). Averaging these two estimates we arrive at a total number of expected ciliary genes of approximately 1200 genes.

### Validation and OSCP1 experiments

#### Candidate selection

The number of genes tested per method, the model organisms and cell-lines were determined by time and resources available to the participating labs. Orthologs in *C. elegans* were identified for the candidate list. For 21 orthologs null alleles were available from the *Caenorhabditis* Genetics Center (University of Minnesota, USA). All 21 null mutants were obtained and tested. Candidates for the eCFP localization studies in hTERT-RPE1 cells were randomly selected. The candidates for the immunofluorescence in human lung epithelial and murine retina were selected based on the availability of a suitable antibody in the collaborating labs. Candidates tested in zebrafish were randomly selected based on the presence of unambiguous 1-1 orthologs. An equal spread of the selected candidates throughout the ranked list was taken into account as much as possible as is shown in Fig. 4h.

#### Localization studies in hTERT-RPE1 cells

Expression constructs were created with Gateway Technology (Life Technologies) according to the manufacturer’s instructions. These constructs encoded eCFP fusion proteins of OSCP1 (transcript variant 1; NM_145047.4), CFAP61 (C20orf26, NM_015585), CFAP20 (C16orf80, NM_013242.2), CYB5D1 (NM_144607), CCDC147 (M_001008723.1), CFAP161 (C15orf26, NM_173528), IQCA1 (NM_024726.4), IPO5 (NM_002271), HSPA1L (NM_005527), and TEKT1 (NM_053285). The sequences for all entry clones were verified by Sanger sequencing. Human TERT-immortalized retinal pigment epithelium 1 (hTERT-RPE1) cells were cultured as previously described^82^. Cells were seeded on coverslips, grown to 80% confluency, and subsequently serum starved for 24 hr in medium containing only 0.2% foetal calf serum for inducing cilium growth. The cells were then transfected with eCFP expression construct using Lipofectamine 2000 (Life Technologies) according to the manufacturer’s instructions. Cells were fixed in 4% paraformaldehyde for 20 min, treated with 1% Triton X-100 in PBS for 5 min, and blocked in 2% BSA in PBS for 20 min. Cells were incubated with the primary antibody GT335 (cilium and basal body marker, 1:500) diluted in 2% BSA in PBS, for 1 hr. After washing in PBS, the cells were incubated with the secondary antibody for 45 min. Secondary antibody, goat anti-mouse Alexa 568 (1:500; Life Technologies) was diluted in 2% BSA in PBS. Cells were washed with PBS and briefly with milliQ before being mounted in Vectashield containing DAPI (Vector Laboratories). The cellular localization of eCFP-fused proteins was analyzed with a Zeiss Axio Imager Z1 fluorescence microscope equipped with a 63x objective lens. Optical sections were generated through structured illumination by the insertion of an ApoTome slider into the illumination path and subsequent processing with AxioVision (Zeiss) and Photoshop CS6 (Adobe Systems) software.

#### Immunofluorescence microscopy of cells

IMCD3 and ATDC5 cells were growth to 80% confluence in DMEM-Glutamax medium with 10% Foetal Bovine Serum. Then cells were Serum-starved for 24 hours and fixed in cold methanol for 5 minutes, PBS washed and blocked with 1% Bovine Serum Albumin for 1 hour before incubating with primary antibodies overnight at room temperature. Antibodies and concentrations were anti-acetylated -tubulin (Sigma 6-11B-1, T7451) 1/200, anti-gamma-tubulin (Sigma GTU-88, T6557) 1/200, anti-OSCP1 Proteintech 12598-1-AP 1/100, anti-OSCP1 ATLAS HPA028436 1/100 and anti-OSCP1 Biorbyt 185681 1/100. Human respiratory cells were analyzed by immunofluorescence microscopy as previously described^83^. The following rabbit polyclonal antibodies were purchased from Atlas antibodies: anti-CCDC113 (HPA040869), anti-RIBC2 (HPA003210), anti-ARMC3 (HPA037824), anti-C6orf165 (HPA044891) and anti-OSCP1 (HPA028436).

#### Mouse handling and experiments

C57Bl/6J wild-type mice were kept on a 12 h light-dark cycle with unlimited access to food and water. All procedures were in accordance with the guidelines set by the ARVO statement for the use of animals in Ophthalmic and Vision Research and the local laws on animal protection. The following antibodies were used for immunofluorescence/immune-EM analysis of murine retina sections: rabbit anti-CCDC113 (1:500/1:500), anti-RIBC2 (1:250/1:250), anti-EFHC1 (1:500/-), anti-WDR69 (1:250/-), C6orf165 (1:50/1:200), anti-ARMC3 (1:250/1:500), anti-OSCP1 (for IF 1:100, biorbyt, Cambridge, UK; 1:100, proteintech, Manchester, UK; for EM: 1:100, Atlas, Stockholm, Sweden), mouse anti-centrin-3 (1:100;^84^), rabbit anti-rootletin (1:100, ^85^). Sections were counterstained with DAPI (1 mg/ml) Sigma-Aldrich, Munich, Germany), and, where applicable, with FITC-labeled peanut agglutinin (PNA, 1:400, Sigma-Aldrich, Munich, Germany). Secondary antibodies conjugated to Alexa 488^®^, Alexa 555^®^, and Alexa 568^®^ (1:400) were purchased from Invitrogen (Karlsruhe, Germany) and CF™-640 (1:400) from Biotrend Chemikalien GmbH (Cologne, Germany). For pre-embedding electron microscopy, we used biotinylated secondary antibodies (1:150; Vector Laboratories, Burlingame, CA, USA). For immunofluorescence microscopy, the eyes of adult C57Bl/6J mice were cryofixed sectioned, and immunostained as described previously ^86^}. We double-stained the cryosections for CCDC113, RIBC2, EFHC1, WDR69, C6orf165, ARMC3, OSCP1, and centrin-3 as a molecular marker for the connecting cilium, the basal body, and the adjacent centriole of photoreceptor cells {Trojan et al. 2008} at 4°C overnight. Sections stained for C6orf165 were counterstained with FITC-labeled cone photoreceptor marker peanut agglutinin (PNA^87-89^). After one-hour incubation at room temperature with the according secondary antibodies and the nuclear marker DAPI, sections were mounted in Mowiol 4.88 (Hoechst, Frankfurt, Germany). Images were obtained and deconvoluted with a Leica LEITZ DM6000B microscope (Leica, Wetzlar, Germany) and processed with Adobe Photoshop CS with respect to contrast and color correction as well as bicubic pixel interpolation. We applied a pre-embedding labeling protocol as previously introduced for immunoelectron microscopy of mouse photoreceptor cells ^90-92^. Ultrathin sections were analyzed with a transmission electron microscope (TEM) (Tecnai 12 BioTwin; FEI, Eindhoven, The Netherlands). Images were obtained with a charge-coupled device camera (SIS MegaView3, Olympus, Shinjuka, Japan) and processed with Adobe Photoshop CS (brightness and contrast).

#### Zebrafish handling and experiments

Wild-type (AB × Tup LF) zebrafish were maintained and staged as described previously in^93^. Antisense MO oligonucleotides (Gene Tools) were designed against the start codons and against splice sites, as described in Supplementary table 6. MOs were injected (4-6 ng) into embryos at the 1- to 2-cell stage and reared at 28.5°C until the desired stage. For cilia immunostaining, 6 somite-stage or 24 hpf (hours post fertilization) embryos were dechorionated and fixed in 4% PFA overnight (O/N) at 4°C, dehydrated through 25%, 50% and 75%, methanol/PBT (1% Triton X-100 in PBS) washes and stored in 100% methanol -20°C. The embryos were rehydrated again through 75%, 50% and 25% methanol/PBT washes. Embryos of 24 hpf were permeabilized with Proteinase K (10ug/ml in PBT) for 10 minutes at 37°C, and subsequently refixated in 4% PFA. Prior to immunostaining, embryos were incubated in block buffer (5% goat serum in PBT) blocked with 5% goat serum (in PBT) for 1 h and subsequently incubated O/N at 4oC with mouse monoclonal anti-γ-tubulin (1:200, GTU-88, Sigma) and anti-acetylated tubulin (1:800, 6-11B-1, Sigma) diluted in blocking buffer. Secondary antibodies used were Alexa Fluor goat anti-mouse IgG1 488, Alexa Fluor donkey anti-mouse IgG2b 568, and Alexa Fluor goat anti-mouse IgG2b 594 (Molecular Probes). Nuclei were stained with Hoechst and embryos were mounted in Citofluor. Z-stack images were captured using a Zeiss 710 Confocal Microscope. For rescue experiments, the aforementioned human OSCP1 cDNA was cloned into pCS2+ using gateway technology. OSCP1 plasmids were linearized using NotI and mRNA was synthesized using Ambion mMessage mMachine kit for the sense strand. 100 pg of mRNA was injected into the cell of one cell-stage embryos. These embryos were subsequently injected with 4 ng oscp1 MO at the two-cell stage, and embryos were allowed to develop at 28.5°C.

#### C. elegans handling and experiments

*C. elegans* were maintained and cultured at 20°C using standard techniques. Mutant strains were obtained from the *Caenorhabditis* Genetics Center (University of Minnesota, USA); the alleles used are shown in Supplementary table2. Assays for dye uptake (DiI), roaming and osmotic avoidance were performed as previously described^23^. Briefly, for the dye-filling assay, worms were placed into a DiI solution (diluted 1:200 with M9 buffer) for 1 hour, allowed to recover on NGM plates, and then imaged (40x objective, Texas Red filter set) on a compound epi-fluorescence microscope (Leica DM5000b), fitted with an Andor EMCCD camera. For the osmotic avoidance assay, young adult worms were placed within a ring-shaped barrier of 8M glycerol and scored during 10 minutes for worms that crossed the barrier. For the roaming assay, single young adult worms were placed for 1 hours onto seeded plates and track coverage assessed using a grid reference. For transmission electron microscopy, *oscp-1*(*gk699*) worms were first backcrossed 2 times with wild type worms (to remove unlinked mutations), using primers that flank the *gk699* deletion. Day 1 adults were fixed, sectioned and imaged as described previously^23^. Translational reporters were introduced into*oscp-1*(*gk699*) by standard mating methods. The translational construct for *oscp-1* (R10F2.5 and R10F2.4) was generated by fusing the genomic region, including 853 bp of the native promoter, to GFP with the *unc-54* 3’ UTR. Standard mating procedures were used to introduce OSCP-1::GFP into the *mks-5*(*tm3100*) mutant background.

## Acknowledgements

We kindly acknowledge Prof. Dr. N. Katsanis and J. R. Willer from Duke University USA for providing several of the clone constructs used as baits in the SILAC and Y2H data sets. We also thank the *Caenorhabditis elegans* Genetics Center(Minnesota, USA), the National Bioresource project (Tokyo, Japan), the International *C. elegans* gene knockout consortium, and the *C. elegans*Million Mutation Project for nematode reagents. We thank Emine Bolat, Ideke Lamers, Moniek Riemersma, and Karlien Coene for performing yeast two-hybrid and SILAC experiments. This work was financially supported via the European Community’s Seventh Framework Programme FP7/2009 (SYSCILIA grant agreement no: 241955). TJPvD, RvdL and MAH acknowledge support by the Virgo consortium, funded by the Dutch government (FES0908), and by the Netherlands Genomics Initiative (050-060-452). MRL acknowledges research funding from the Canadian Institutes of Health Research (CIHR; grants MOP-142243 and M0P-82870) and a senior scholar award from Michael Smith Foundation for Health Research (MSFHR). VLJ acknowledges postdoctoral fellowships from MSFHR and KRESCENT. RHG, PLB and RR acknowledge the Dutch Kidney Foundation “KOUNCIL” consortium CP11.18. M.U. and K.B. acknowledge FP7 grant agreement no. 278568, PRIMES. RBR and QL acknowledge support by the Excellence grant CellNetworks funded by the Deutsche Forschungsgemeinschaft (DFG). RS acknowledges support by Metakids Foundation and European Union 7th Framework Programme [PROPANE Study, 602273]. PLB and VH-H acknowledge support by the NIHR Great Ormond Street Hospital Biomedical Research Center. PLB is an NIHR Senior Investigator. MS acknowledges support by a Radboudumc Hypatia Tenure Track Fellowship, a Radboud University Excellence Fellowship and the German Research Foundation (DFG) (collaborative research center grant SFB-1411 KIDGEM). RR acknowledges support by the Netherlands Organization for Scientific Research (NWO Vici-865.12.005).

## Authors’ contributions

TJPvD and MAH conceived and led the study. TJPvdD and MAH, wrote the manuscript with significant contributions from RBR, OEB, RHG, and RvdL. RvdL and TJPvD developed the algorithm and performed the bioinformatic analyses. JK, EdV, KAW, SR, GWD, NJL, CL, VLJ, RH, and VH-H performed the validation experiments and experiments on OSCP1. EdV, NH, YT, YW, JR, GW, BK, JFS, DAM, EvW, GGS, KS, TMTN, SJFL, SECvB, and KB performed experiments leading to the Y2H and SILAC data sets. TJPvD, RvdL and RS created the bioinformatics data sets. TJPvD, JvR, KK, GT, QL collected, quality assessed, formatted and mapped the data sets. MRL, FK, HK, HO, MU, PLB, BF, MS, RHG, RBR, TJG, CAJ, OEB, UW, KB, RR, VH-H, GWD, and MAH suggested strategies and supervised work. All authors read and approved the final manuscript.

## Materials & Correspondence

Material requests and correspondence should be addressed to T.J.P. van Dam or Martijn A. Huynen.

